# Striatal dopamine signals internal states related to behavioral performance over prolonged timescales in monkeys

**DOI:** 10.64898/2026.06.24.733789

**Authors:** Raymond Murray, Usamma Amjad, Ann M. Graybiel, James P. Herman, Helen N. Schwerdt

## Abstract

Slowly varying internal states govern our ability to sustain goal-directed behavior over minutes to hours, imposing fundamental constraints on cognitive performance, yet the neural signals that track these states remain poorly defined. Dopamine is a powerful modulator of motivated behavior, but its best-characterized signals are in the form of reward prediction errors (RPEs) that operate at seconds timescales. Whether these fast signals also carry information about slower, ongoing states of behavioral performance has not been directly tested. Here, we recorded subsecond dopamine concentration changes across multiple sites in the caudate nucleus (CN) and putamen of rhesus monkeys performing a reward-guided saccade task. Task performance oscillated over tens to hundreds of trials, demonstrating fluctuating internal states of sustained engagement. We found that single-trial dopamine signals were strongly modulated by these slowly evolving performance states, in a manner dissociable from both trial-level RPE and cumulative reward rate. This relationship was biased toward future rather than past performance windows, indicating that dopamine tracks motivational state prospectively rather than simply reflecting the reward history on which prediction errors are computed. Furthermore, performance-state modulation was concentrated in the CN rather than the putamen, consistent with the distinct roles of these regions in oculomotor and skeletomotor control and suggesting that state-dependent dopamine signals are organized according to the behavioral demands of the task. These findings demonstrate that phasic dopamine signals in the dorsal striatum simultaneously reflect fast learning signals and slowly evolving internal states, linking dopamine’s canonical RPE function to its broader role in sustaining motivated performance.

## Introduction

Sustained cognitive performance varies over timescales of minutes to hours. These performance fluctuations — observed as attention waxing and waning, clustered error rates, and lapses in engagement^1–3^ — are thought to reflect slowly evolving internal states that modulate how reliably actions are initiated and maintained in pursuit of a goal^4–10^. Multiple neural systems^11–16^ have been implicated in regulating such motivational states, but the specific signals that covary with naturally occurring performance fluctuations remain poorly defined.

Dopamine is a powerful modulator of motivated behavior and learning^4,9,17,18^, but its phasic signals are best characterized at the timescale of individual trials (seconds). These neurotransmitters are known to signal reward prediction errors (RPEs) and related reward-value computations^17,19–25^: an unexpected reward produces a phasic dopamine increase but a fully predicted reward does not. This creates a puzzle: dopamine is widely implicated in regulating sustained motivational states, yet its best-characterized signals are transient and trial-bound. Whether these transient signals carry information about ongoing internal states that shape behavioral performance over minutes to hours has not been directly tested.

Evidence linking dopamine to motivational states over longer timescales has grown — from classic pharmacological work showing that dopamine levels bidirectionally control willingness to work for reward^4,9^, to more recent demonstrations that dopamine concentration fluctuations track reward rate^5^ and are modulated by homeostatic drives such as hunger and thirst^26^. However, these longer-timescale links have been attributed to slower dopamine dynamics that are dissociable from the single-trial phasic transients that encode RPE^27^. Whether the same phasic dopamine responses that carry trial-level reward signals can also carry information about slower internal states of behavioral performance remains an open question.

The primate striatum is organized into functionally distinct subregions, the caudate nucleus (CN) and the putamen, which receive distinct cortical inputs and serve separable roles in oculomotor-cognitive and skeletomotor control^28–31^. Dopamine innervation is topographically organized across these structures, and recent work has shown that dopamine signaling reflects this functional topography in its representation of reward-related variables^27,32–34^. Mechanisms for locally modulating dopamine release at striatal terminals have also been identified^35,36^, raising the possibility that dopamine signals are shaped by the distinct behavioral demands of each region. If dopamine signals related to performance state exist in dorsal striatum, they may differ between the CN and putamen.

We recorded dopamine concentration changes across multiple (37) chronically implanted sites in the CN and putamen of two rhesus monkeys using established fast-scan cyclic voltammetry (FSCV) protocols^37–39^. Monkeys performed a reward-biased visually-guided saccade task over hundreds of trials per session, producing fast (∼seconds) dopamine signals associated with individual trial events embedded within slower (∼minutes to hours) fluctuations in behavioral performance. We asked: (1) whether single-trial dopamine signals are associated with prolonged internal states of cognitive performance; (2) whether this relationship holds temporal structure, such as a bias toward past or future performance windows; and (3) whether state-dependent dopamine responses differ between CN and putamen. We found that single-trial dopamine release tracks slower states of ongoing task performance in a prospective manner, dissociable from RPE, reward rate, and peripheral autonomic proxies of arousal. This state-dependent relationship was concentrated in the CN, consistent with its role in the oculomotor demands of the task, and provides evidence that phasic dopamine signals in the dorsal striatum encode internal states that sustain motivated performance.

## Results

### Multi-site dopamine recordings in the caudate nucleus (CN) and putamen

Dopamine measurements were made using arrays of carbon fiber (CF) electrodes implanted in two rhesus monkeys, targeting 37 sites in the striatum (**Fig. 1b**)^32,40,41^. Fast-scan cyclic voltammetry (FSCV) was used to measure rapidly fluctuating dopamine concentration changes (Δ[DA]) at millisecond resolution. Sites in the CN were rostral to the anterior commissure (AC) in both monkeys; sites in the putamen were rostral to the AC in monkey P, but were mostly caudal in monkey T. All implanted sensors were maintained chronically, allowing measurements to be reproduced across days^32,41^. Dopamine measurements are reported as area-under-curve (AUC) metrics for Δ[DA]. Results are presented both pooled across sessions for each site and at the level of individual site-sessions (each site treated separately for each session), as previously described^32^.

**Figure 1.**
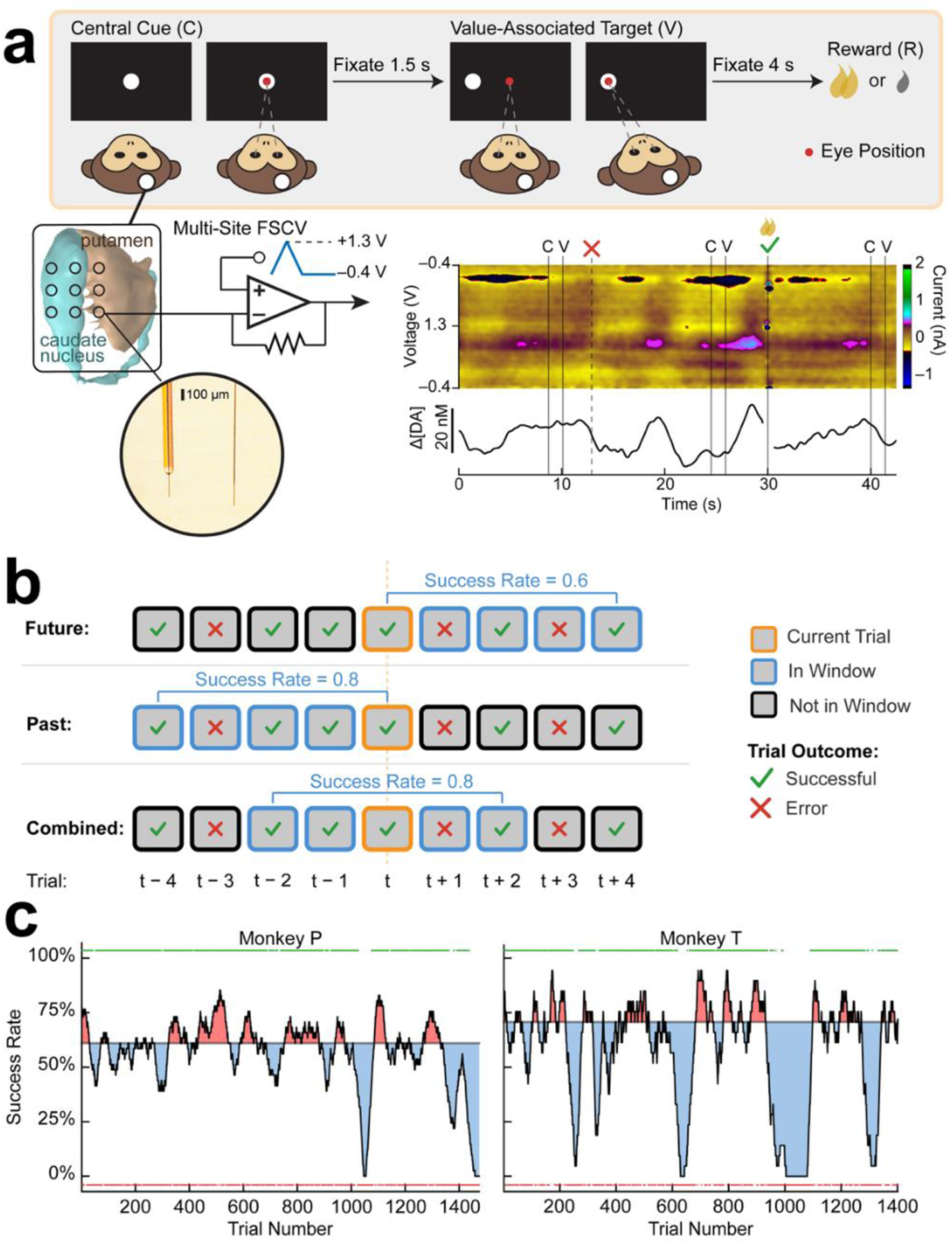
Task design, recording setup, and performance metric. (**a**) Dopamine recordings while monkeys performed a visually-guided reward-biased task (top panel). Target location of the second, peripheral target (V) was associated with reward size (large, yellow, or small, gray) available to the monkey after successful trial completion. Target size-side contingencies were reversed in a block structure (15 – 45 trials for monkey P and 45 – 75 for monkey T). Custom-designed carbon fiber electrodes (photo inset) for FSCV recording of dopamine (see **Methods**) were inserted into the striatum of two female rhesus monkeys using a grid layout that was co-registered with MRI for estimating target trajectories. Color plot shows an example FSCV recording with visible current (color) changes at the redox potentials for dopamine (∼ 0.6 V and –0.2 V) while the monkey performed the task showing an example error trial followed by a successful trial. The extracted dopamine concentration change (Δ[DA]) is shown below it. Photo of the carbon fiber electrodes was reused with permission from a prior publication^32^. (**b**) Moving average success rate metric. Performance was quantified as the number of successful trials divided by the number of trials in the window computed using three windowing methods: future-looking performance (top), backward-looking performance (bottom), and a combined window (middle). (**c**) Moving average success rate (black) as a function of trial number. Using our moving average windowed success rate metric, animal performance was quantified at a trial-by-trial level (monkey P: session 83, combined window, window size 40; monkey T: 3/18/2025, combined window, window size 20). Performance state classification (“high”: red shading, “low”: blue shading) was determined by a trial’s windowed success rate in relation to a success rate threshold, which was defined as the median windowed success rate of successful trials (gray horizontal line). Green dots above the plotted success rates indicate success on the corresponding trial while red dots indicate error.

### Reward-biased saccade task

Both monkeys performed a reward-biased visually-guided saccade task over hundreds of trials per session^28–30^ (**Fig. 1a**). On each trial, the monkey fixated a central cue, then made a saccade to a peripheral target and maintained fixation (3.5 s for monkey T, 4 s for monkey P) to obtain a liquid reward. The reward magnitude (liquid volume) was contingent on the direction of the target (left/right in both monkeys, or up/down in monkey T), and this contingency was maintained for a block of trials (45 – 75 in monkey T, 15 – 45 in monkey P) before being reversed. Monkeys routinely completed more than a thousand trials per session, and success rates, as calculated across a window (10 – 100) of trials, varied both across days and within sessions (**Fig. 1c**).

All dopamine signals and physiological measures analyzed in this study were taken from the target fixation period of each successful trial, unless otherwise noted. The target cue indicated the upcoming reward size; this is known to consistently evoke dopamine signaling activity^5,17,42^. Saccade latencies confirmed that reward expectation was behaviorally expressed at the target cue: latencies tracked the cue’s location-reward contingency reliably, except on the first trial after a block switch (**Extended Data Figure 1**). Latencies were not updated until the following trial, consistent with the monkey being unaware of the reversal until after experiencing the outcome.

### Expected reward size modulates behavioral and autonomic measures at single-trial timescales

Expected reward size modulated multiple behavioral and autonomic measures at the single-trial level (**Figure 2**). Movements were faster for higher value target cues with saccade latencies being shorter for large-reward targets than small-reward targets in both monkeys across all sessions. Anticipatory licking also scaled with expected reward (12/15 sessions in monkey P, 8/9 sessions in monkey T) indicating higher conditioned reward responding and motivation^9,43^. Pupil diameter, a proxy for arousal, cognitive load, and general salience^11,44^, was more dilated for large-reward than small-reward trials (14/15 sessions in monkey P, 5/9 sessions in monkey T; two-sided standard permutation test, p < 0.05). Heart rate variability (HRV), an indirect marker of parasympathetic activity and vagal output^45,46^, was also modulated by reward size (13/15 sessions in monkey P, 5/9 sessions in monkey T). The direction of this HRV modulation varied by session.

**Figure 2.**
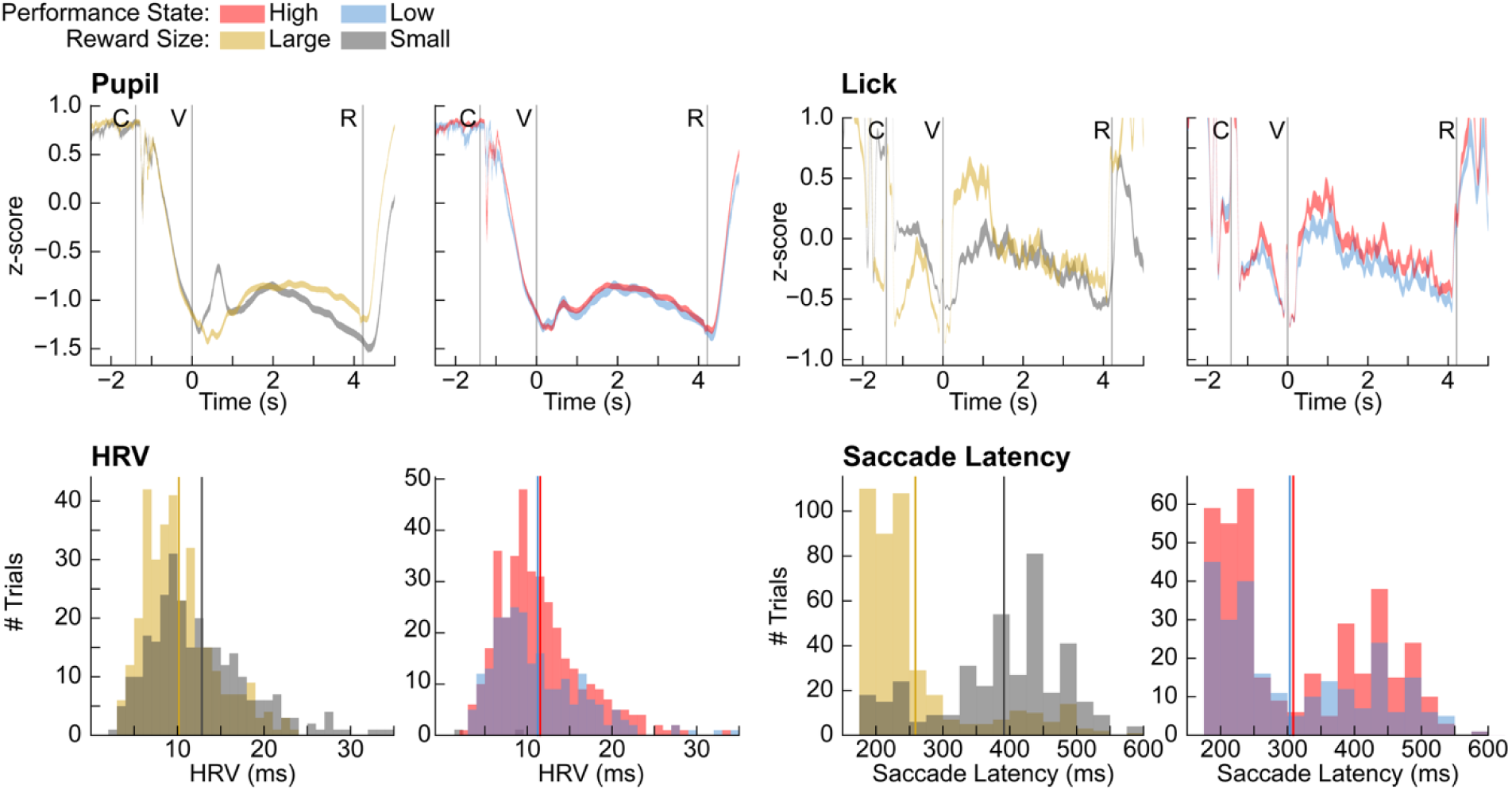
Non-neural physiological responses differ by reward magnitude. Examples of concurrently measured non-neural physiological responses split by reward condition (left for each grouping) and performance state (right) from monkey P. Pupil diameter (z-scored; top left; session 102), anticipatory licking (z-scored; top right; session 84), heart rate variability (HRV; bottom left; session 114), and saccade duration (bottom right; session 95) are shown.

### Performance states defined by fluctuating success rates

Task performance fluctuated markedly within a session, with success rates oscillating over windows of tens to hundreds of trials (**Fig. 1c**). We captured these fluctuations by classifying each trial as occurring during a high or low performance state based on whether the windowed success rate exceeded the session median (**Fig. 1b**). The window was varied systematically from 10 to 100 trials rather than being fixed at a single value to evaluate the timescales over which performance states showed dopamine modulation (**Extended Data Figure 2**). An extended kernel regression analysis presented in later sections provides a complementary continuous approach that does not depend on this discretization. This classification allowed direct comparison of single-trial dopamine dynamics with the slower performance state in which each trial was embedded.

Behavioral and autonomic measures were only weakly and inconsistently associated with performance state (**Figure 2**), contrasting sharply with their robust reward-size modulation. Lick amplitude and saccade latencies were each modulated by performance state in fewer than half of sessions in either monkey (lick: 4/9 monkey T, 3/15 monkey P; saccade latency: 4/9 monkey T, 4/15 monkey P), and when saccade latency effects were present, their direction was inconsistent. Pupil diameter and HRV reached significance in more sessions but with no consistent direction across monkeys (pupil: 6/9 monkey T, 8/15 monkey P; HRV: 6/9 monkey T, 9/15 monkey P). The same behavioral and autonomic signals that reliably differentiated reward conditions thus failed to consistently differentiate performance states, and performance state could not be reduced to any single observable behavioral or autonomic proxy^47^.

### Cue evoked dopamine responses differentially reflect reward value and RPE on individual trials

Phasic dopamine has been shown to encode reward value and prediction error at the timescale of individual trials^17,48–52^. We sought to confirm whether our chronic FSCV measurements also captured these canonical signals by examining single-trial dopamine as modulated by expected reward size and RPE. Nearly half of recorded sites displayed significant reward-size modulation, when pooled across sessions (8/16 CN sites and 9/21 putamen sites; **Figure 3**). Consistent with value coding, dopamine release was greater for large-reward than small-reward at these reward modulated sites, except for one site in the putamen in monkey P.

**Figure 3.**
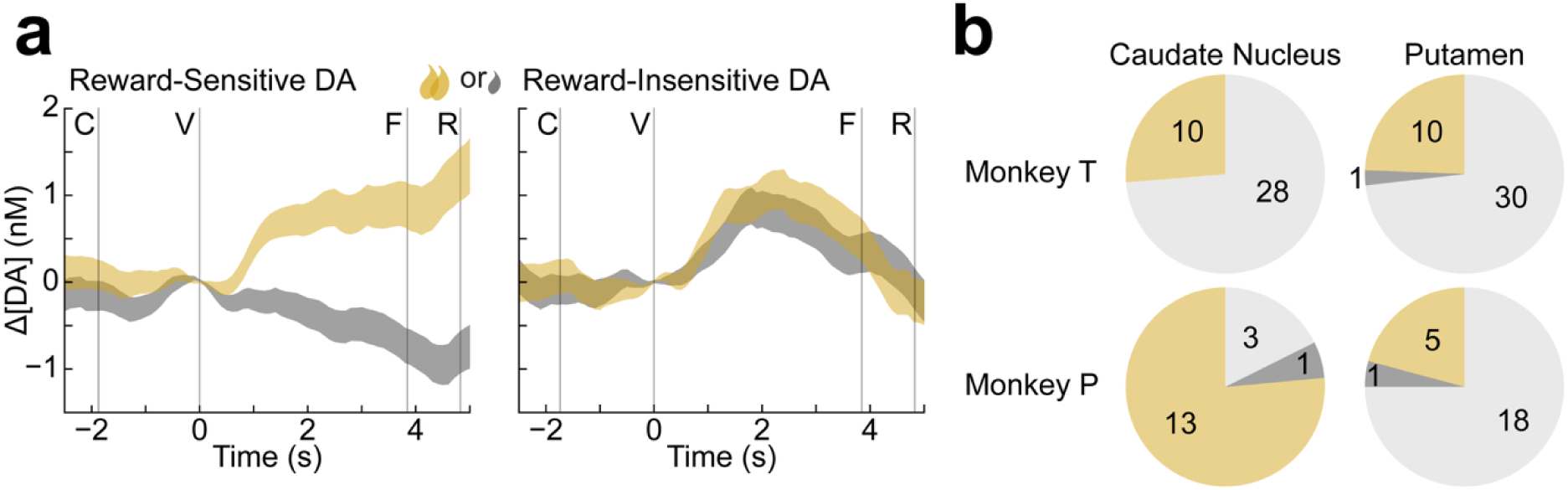
Single-trial dopamine release differs by reward magnitude. (**a**) Trial-averages for dopamine changes measured in monkey T denoting examples where dopamine was sensitive to reward size (left; c5c recorded on 9/24/2024) and insensitive to reward size (right; c8c recorded on 10/3/2024). Shading indicates ±SEM. Vertical lines denote task events: central cue onset (C), peripheral cue onset (V), visual trial-completion feedback (F, monkey T only), and reward delivery (R). (**b**) Site-session counts of reward-sensitive dopamine in the CN (left) and putamen (right) for monkeys T (top) and P (bottom; two-sided standard permutation test, *p* < 0.05). Coloring denotes which reward size showed larger DA AUCs (large: yellow, small: dark gray) or no difference (light gray).

RPE was estimated using a Rescorla-Wagner model in which value estimates were updated at the peripheral target cue rather than at reward delivery (see **Methods**). This cue- RPE formulation follows from temporal difference learning, in which the prediction error signal transfers to the earliest reliable reward predictor^17,19^. In our task, the target cue indicates the upcoming reward size and is therefore the earliest predictor. Saccade latencies confirmed that reward expectations were behaviorally expressed at this cue, with faster movements being made for larger upcoming rewards. These reward-contingent behaviors also updated immediately (within a single trial) following block reversals (**Extended Data Figure 1**). Cue-evoked dopamine correlated significantly with RPE in both monkeys (CN: 5/7 and 6/9 sites; putamen: 5/10 and 6/11 sites in monkeys T and P, respectively) (**Figure 4**). Collectively, the reward-modulated and RPE-correlated dopamine responses are consistent with canonical theories of dopamine’s role as a teaching signal in reinforcement learning^48–51^.

**Figure 4.**
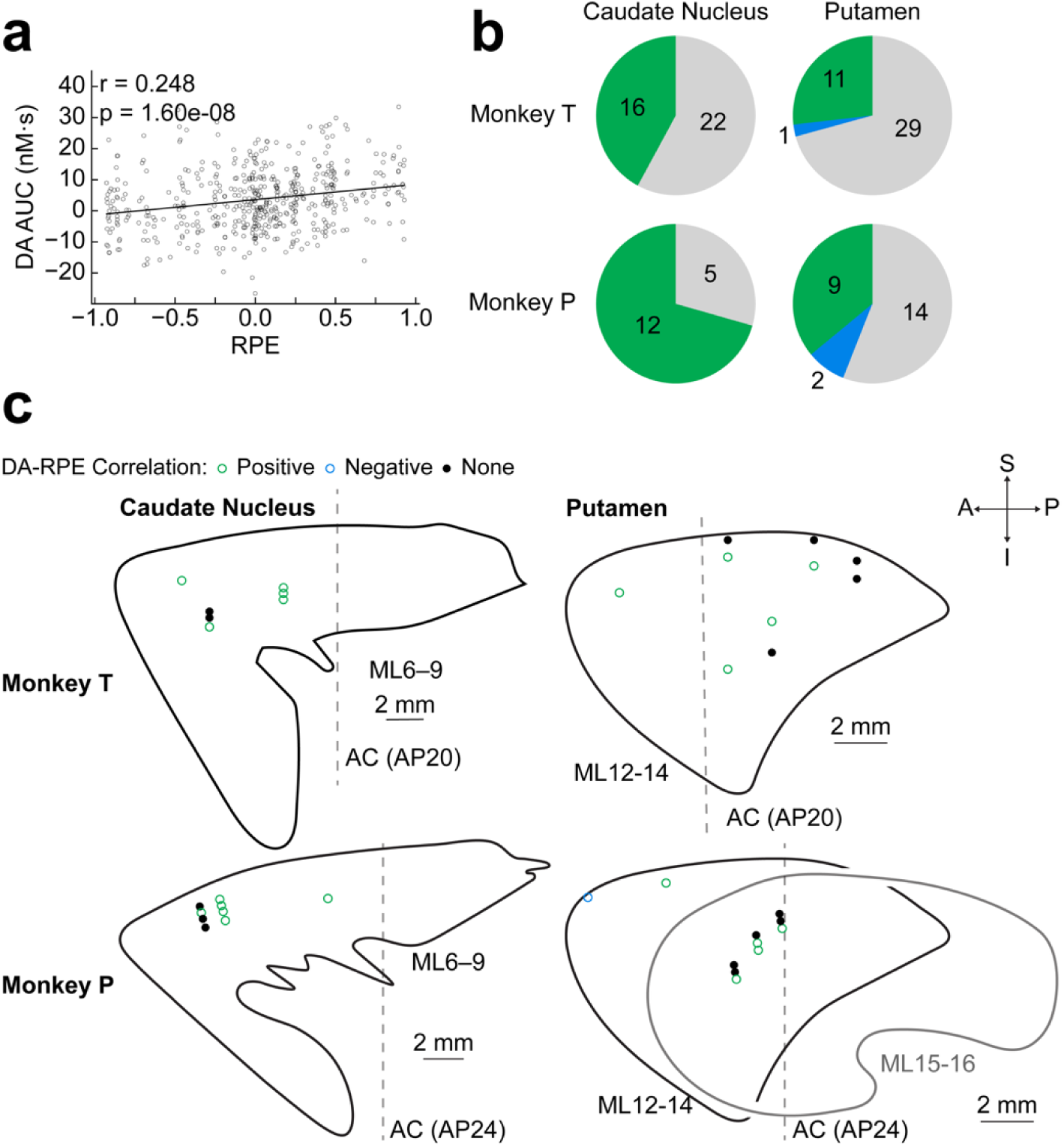
Single-trial dopamine release is correlated to reward prediction error (RPE) in some sites. (**a**) Example correlation between model-derived RPE and dopamine concentration changes, computed as AUC values (DA AUC), in monkey P (c64 recorded on session 102). (**b**) Site-session counts of statistically significant correlations between RPE and DA AUC, color-coded by the sign of the spearman correlation coefficient. (**c**) Sagittal sections showing recording site locations in the caudate nucleus and putamen for monkey T (top) and monkey P (bottom), color-coded by the sign of the spearman correlation coefficient.

### Single-trial dopamine release is strongly modulated by prolonged performance states

We quantified the relationship between single-trial dopamine signaling and prolonged performance states by comparing the signals between trials occurring during high versus low performance states. These performance states were defined using a sliding window of 10 – 100 trials centered on each trial (see **Methods**). Dopamine release was greater during high than low performance states in most of the CN sites in both monkeys (monkey T: 7/7 sites, 24/38 site-sessions; monkey P: 5/9 sites, 9/17 site-sessions; two-sided circular permutation tests, all p < 0.05; **Figure 5**). Modulation of dopamine release levels in the putamen was also present, but was less consistent, with stronger effects in monkey T (8/10 sites, 20/41 site-sessions) than monkey P (2/11 sites, 5/25 site-sessions). A minority of sites, only in monkey P, showed the opposite pattern, with lower dopamine during high performance states; this was more frequent in the putamen of monkey P (3/11 sites) than in the CN (1/9 sites).

**Figure 5.**
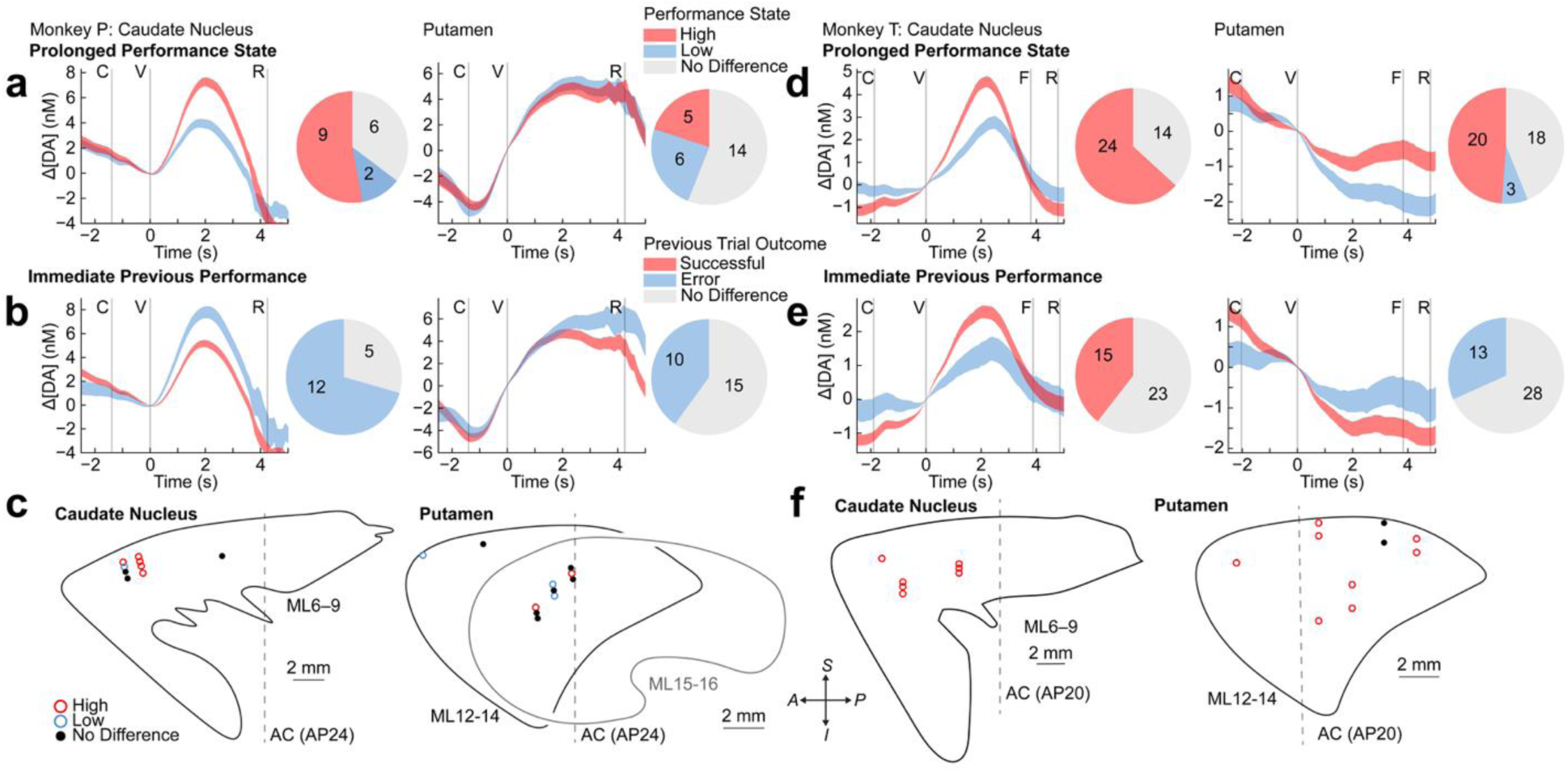
Dopamine reflects prolonged states related to performance in a manner that contrasts with its association with immediate performance outcomes. Trial-averaged striatal dopamine (Δ[DA]) in monkey P split by better (red) vs. worse (blue) performance trials, for long timescale (**a**) and short timescale (**b**) performance classifications (defined as the effect of success or error on the immediate previous trial), in the caudate nucleus (left, c55 recorded on session 94) and putamen (right, p033 recorded on session 67). Shading denotes ±SEM. Pie charts indicate the proportion of site-session counts exhibiting each dopamine-performance scaling pattern (color-coded as in legend; gray = performance-agnostic). (**d–e**) Same as **a–b** for monkey T (caudate nucleus: c8ds recorded on 9/30/2024 for prolonged performance state and 9/25/2024 for immediate previous performance; putamen: p4d recorded on 3/18/2025 for both prolonged performance state and immediate previous performance). (**c,f**) Sagittal sections showing recording site locations in the caudate nucleus and putamen for monkey P (**c**) and monkey T (**f**), color-coded by dopamine-performance state classification under short vs. long timescales. Pie charts denote site-session counts for statistically significant (*p* < 0.05) differences in dopamine AUC distributions as determined by two-sided circular permutation tests for the prolonged performance state and standard permutation tests for immediate previous performance.

The performance-state effect was independent of RPE signaling. Residual analysis in which dopamine was first regressed on trial-by-trial RPE confirmed that the performance-state relationship persisted across the vast majority of site-sessions in both monkeys (65/69 performance-modulated site-sessions; two-sided circular permutation tests, all p < 0.05; **Extended Data Figure 3**).

The prolonged performance-state modulation contrasted with the effect of the immediately preceding trial outcome. Dopamine release tended to be lower following a successful trial than a failed trial. This finding is opposite to the performance-state effect, where dopamine was higher with higher success rate, but consistent with RPE-based updating in which a recent success raises expected value and reduces the prediction error on the next trial (**Extended Data Figure 4**). This pattern was more consistent in the putamen of both monkeys (monkey T: 5/10 sites; monkey P: 6/11 sites; two-sided standard permutation tests, *p* < 0.05) and in the CN of monkey P (8/9 sites). CN sites in monkey T showed the reverse: dopamine was higher following successful trials, a pattern aligned with the performance-state direction rather than RPE direction (6/7 sites). Across both monkeys, performance state was a more consistent modulator of single-trial dopamine than the immediately preceding trial outcome. A kernel regression analysis that does not depend on fixed window size corroborated these findings (see below).

### Performance-state dopamine signals are biased towards future behavioral windows

We used a complementary approach that does not rely on a fixed window size to further corroborate the performance state effects. Trial-by-trial Δ[DA] was regressed onto binary trial outcomes at lags spanning ±100 trials, projected onto 12 raised-cosine basis functions (see **Methods**; **Figure 6**). The resulting kernel weights quantify how strongly outcomes at each lag — past and future — were associated with dopamine release on the current trial. Kernel weights extended substantially across both past and future lags, confirming that single-trial dopamine was not shaped solely by the immediately preceding outcome. CN kernels exhibited consistent structure across both monkeys, with positive weights indicating higher dopamine when surrounding trials were successful. Kernels in the putamen were flatter and less structured.

**Figure 6.**
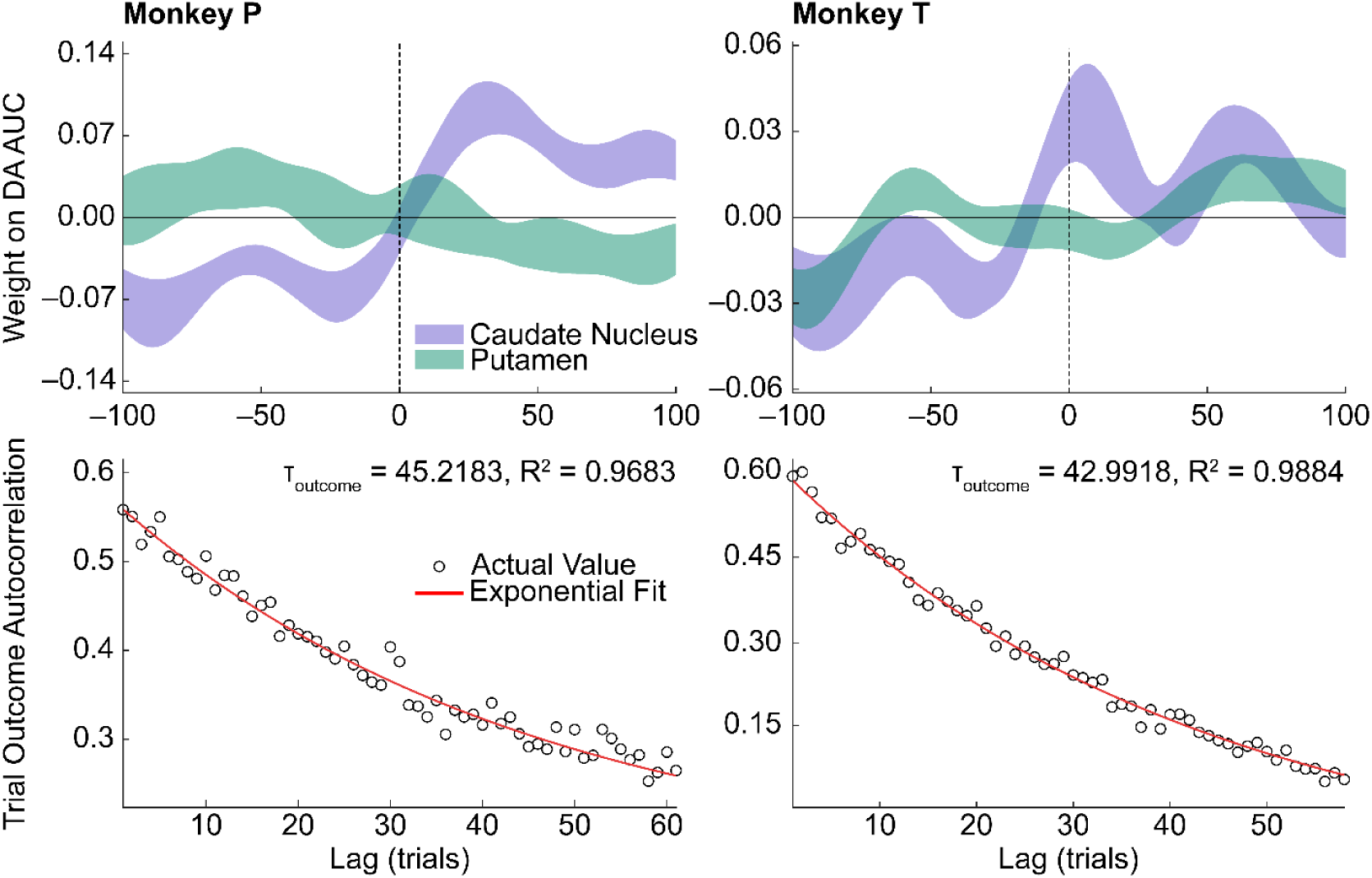
Single-trial dopamine AUC reflects trial outcomes across an extended timescale that matches the autocorrelation on behavior itself. (Top row) Kernel weights from regressing trial-by-trial dopamine AUC onto trial outcomes (binary success/error) at lags spanning ±100 trials for monkey P (left) and monkey T (right). Trial outcomes were projected onto 12 raised-cosine basis functions to smooth across lags and reduce the number of fit parameters (Methods). Shaded bands show mean ± SEM across site-sessions in caudate nucleus (purple) and putamen (green). Dashed vertical line marks lag 0 (current trial); negative lags index past outcomes, positive lags index future outcomes. (Bottom row) Autocorrelation of trial outcomes (open circles) and best-fit exponential decay (red) for representative sessions from monkey P (session 113) and monkey T (3/21/2025). Fitted behavioral timescale parameter τ*_outcome_* (in trials) and goodness-of-fit R² are indicated above each plot.

The kernels were asymmetric: positive weights extended further into future trials than past trials in both regions, indicating that dopamine was more strongly associated with upcoming than recent performance (**Figure 6**). Windowed comparisons corroborated this asymmetry. Single-trial dopamine was more frequently modulated by forward-looking than backward-looking performance windows in both regions (forward: CN 6/7 and 5/9 sites, putamen 6/10 and 4/11 sites in monkeys T and P, respectively; backward: CN 1/7 and 2/9 sites, putamen 3/10 and 4/11 sites; **Figure 7**). The optimal window size for detecting performance-state dopamine differences averaged 54.1 trials in monkey P and 47.5 trials in monkey T for CN sites showing higher dopamine during high performance states (**Extended Data Figure 5**).

**Figure 7.**
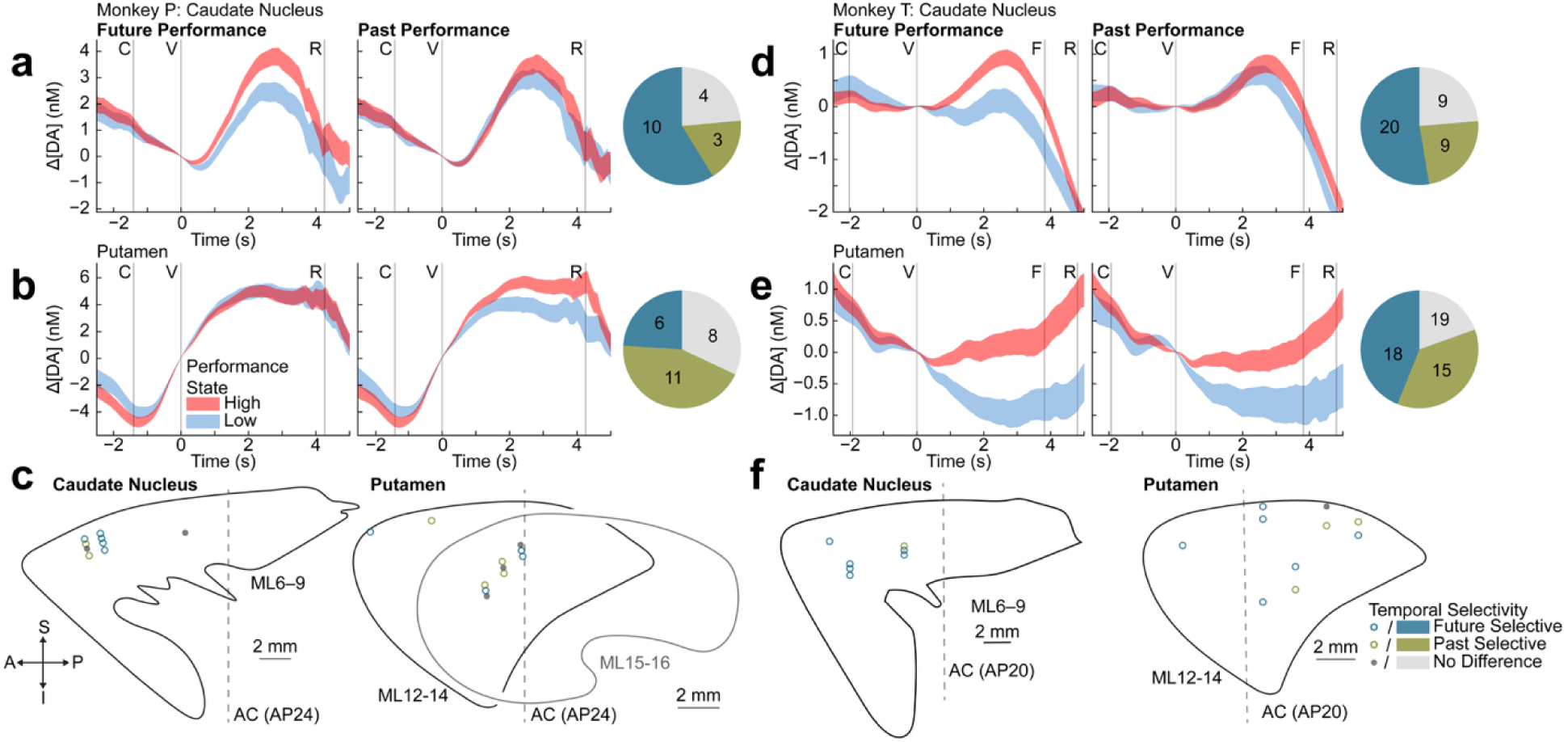
Caudate nucleus dopamine release more reliably tracks future than past behavioral performance across monkeys. (**a, d**) Trial-averaged dopamine concentration changes (Δ[DA]) in the caudate nucleus of monkey P (**a**) (site c54 recorded on session 84) and monkey T (**d**) (c5b recorded on 3/18/2025), split by classified better (red) vs. worse (blue) performance state trials, separately for forward-looking (left) and backward-looking (right) performance classification methods. Shading denotes ±SEM. Pie charts show the proportion of classified as future selective (brown), past selective (orange), or non-selective (gray). (**b, e**) Same as (**a**) and (**d**) for the putamen (monkey P: p033 recorded on session 67, monkey T: p4a recorded on 10/2/2024). (**c, f**) Sagittal sections showing recording site locations in the caudate nucleus and putamen for monkey P (**c**) and monkey T (**f**), color-coded by temporal performance selectivity. Pie charts denote site-session counts where average dopamine AUC differences across performance states were larger when thresholding on future (dark blue) or previous (green) performance.

The temporal structure of behavior itself was consistent with these windowed performance-state dopamine differences. We computed the autocorrelation function (ACF) of trial outcome sequences for each session to independently characterize the timescale of behavioral fluctuations. The ACF measures how strongly outcomes on nearby trials are correlated; a slowly decaying ACF indicates that successes and failures cluster in extended runs rather than occurring independently. Exponential fits to the ACF yielded time constants (τ*_outcome_*) concentrated between 20–60 trials in sessions with significant fits (monkey P: 6/15 sessions; monkey T: 5/9 sessions). This behavioral timescale closely matched the performance- state windows over which dopamine modulation was strongest, providing independent confirmation that the fluctuations captured by our analyses persisted over tens of trials. Together, these results indicate that single-trial dopamine signals were more strongly associated with future than past performance, consistent with a prospective rather than retrospective relationship between dopamine and behavioral state.

### Performance-state dopamine signals are stronger than reward rate modulation, especially in the CN

Higher success rates necessarily produce higher reward rates, raising the question of whether performance-state dopamine differences could be attributed to cumulative reward rate rather than performance state per se. These two rates could be dissociated in our task given the different reward sizes (large and small) available on each trial, in addition to the error trials (no reward). We classified trials as belonging to a high or low reward rate group, similar to the split made for performance states, and compared dopamine release across these two groups. In the CN, performance state was a stronger modulator of single-trial dopamine than reward rate in both monkeys (monkey T: 20/38 vs. 10/38 site-sessions; monkey P: 10/17 vs. 5/17 site-sessions; **Figure 8**). In the putamen, the two measures were comparably associated with dopamine in monkey T (17/41 vs. 17/41 site-sessions), while reward rate was more frequently associated than performance state in monkey P (14/25 vs. 5/25 site-sessions). The stronger dissociation in CN may reflect its established role in oculomotor functions central to our task^28,30^. Performance-state dopamine modulation in the CN was not reducible to cumulative reward history alone.

**Figure 8.**
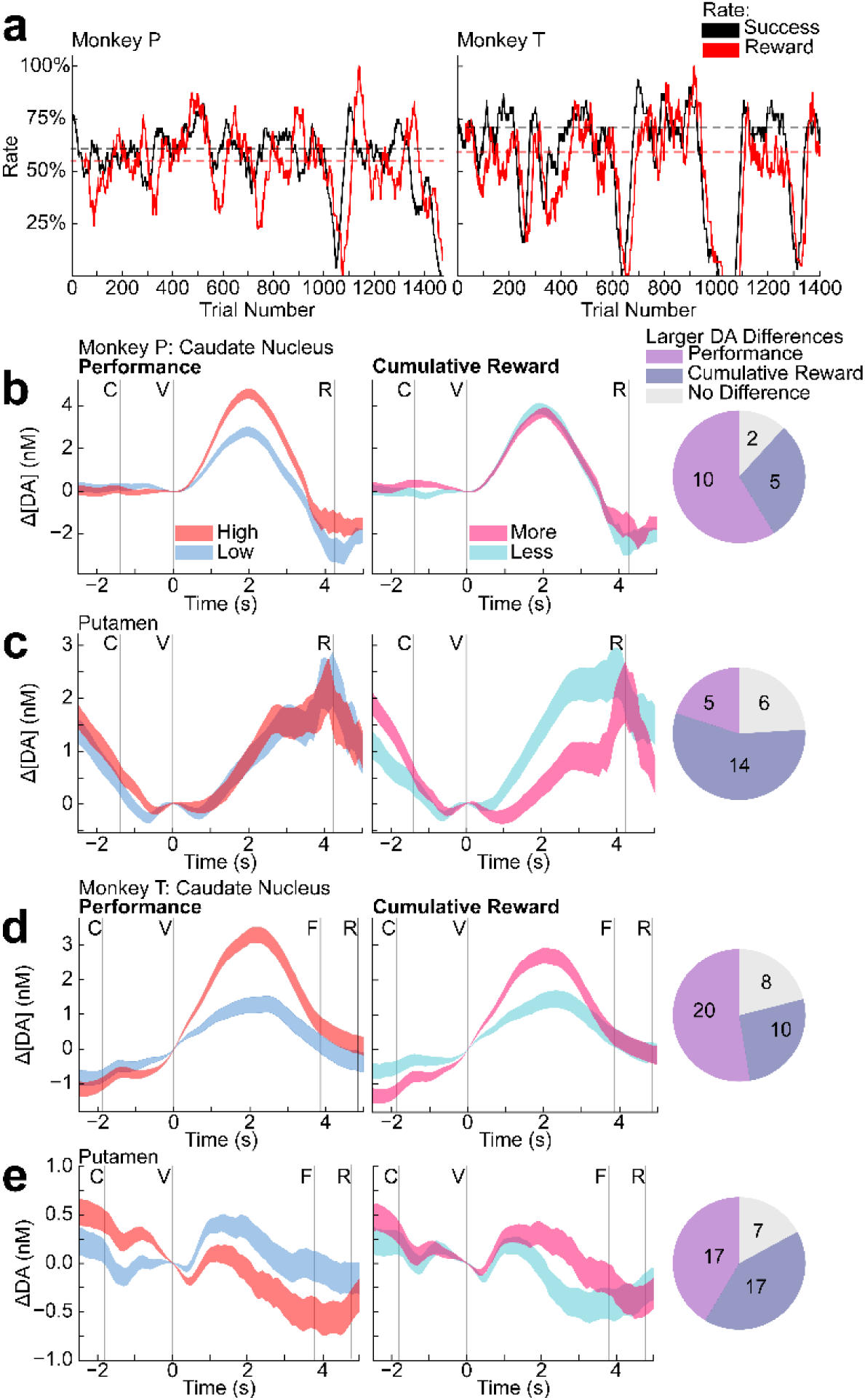
Performance-based trial splits show stronger dopamine differences than cumulative reward in the caudate nucleus of both monkeys. (**a**) Moving average success rate (black) and reward rate (red) across trials for monkey P (left, session 83) and monkey T (right, 3/18/2025). Horizontal dashed lines indicate the median split thresholds used to classify trials as high vs. low performance state (black) or more vs. less rewarded (red). (**b, c**) Trial-averaged dopamine concentration change (Δ[DA]) in the caudate nucleus (**b**, c56 recorded on session 109) and putamen (**c**, p053 recorded on session 102) of monkey P, split by performance (left; better = red, worse = blue) and cumulative reward (right; more = hot pink, less = sky blue). Shading denotes ±SEM. Pie charts show the proportion of site-session counts classified as performance-biased (pink), reward-biased (purple), or non-selective (gray), where bias indicates greater dopamine differences. (**d, e**) Same as (**b, c**) for monkey T (caudate nucleus: c8ds recorded on 9/25/2024; putamen: p2e recorded on 10/3/2024).

### Error type does not affect dopamine’s relation to performance state

Different types of errors were compared to determine whether performance state associations were driven by a particular form of task failure. We separately examined dopamine during periods of elevated total errors, fixation errors (breaks in central or target fixation), and omissions (failure to initiate a saccade). However, this could only be evaluated in monkey T who displayed both omission and commission errors throughout the session, while monkey P rarely committed omissions and these were made only at the end of the recording session. In monkey T, performance-state dopamine differences were present regardless of error type. Dopamine was higher when errors were fewer in all three categories among site-sessions with significant performance-state modulation, in both CN and putamen, (commission: 10/24 CN, 7/23 putamen; omission: 8/24 CN, 8/23 putamen) (**Figure 9**). A single putamen site-session (1/23) showed the opposite pattern for commission errors. This suggests that the performance-state effect was not attributable to any single failure mode.

**Figure 9.**
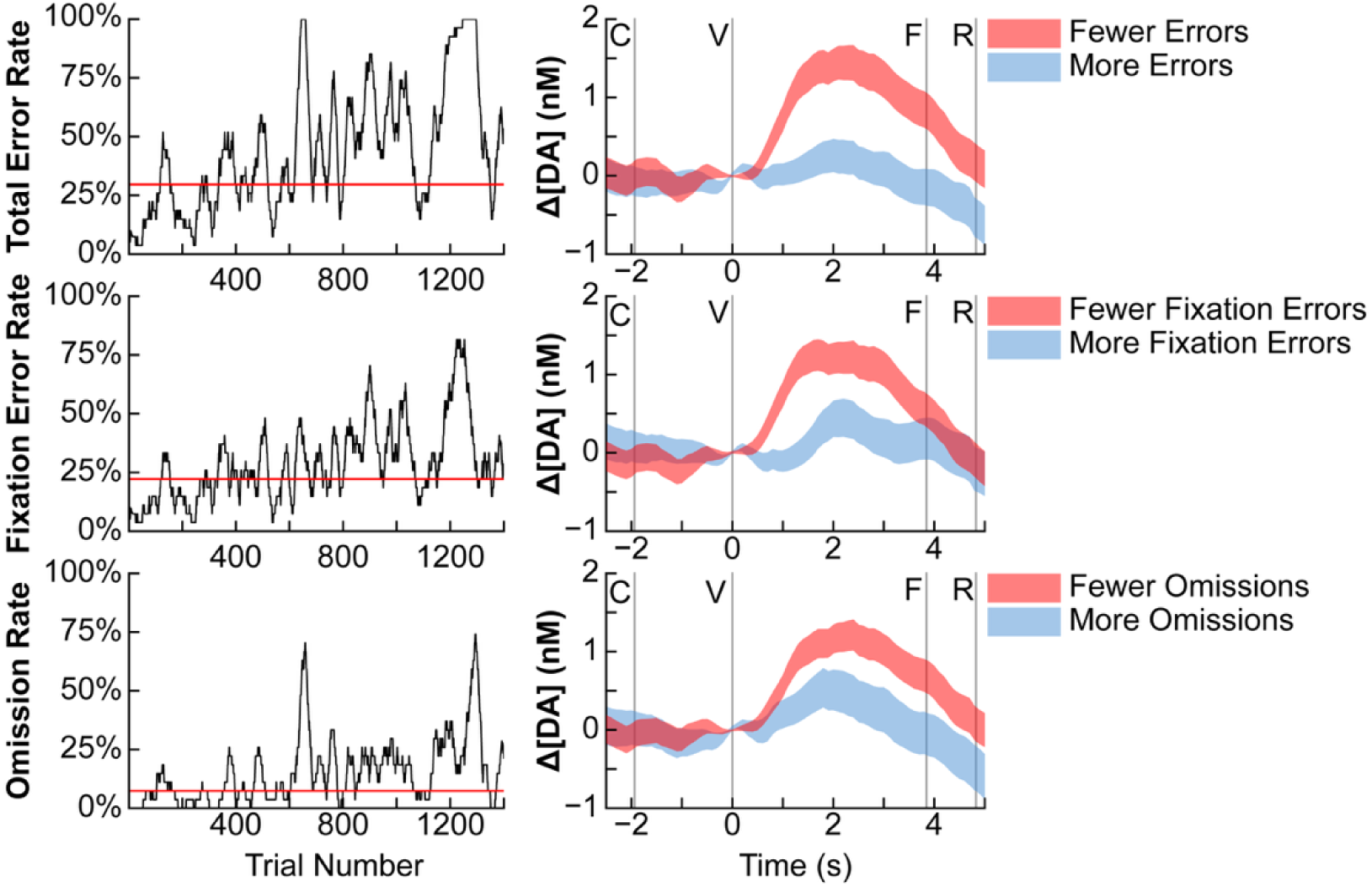
Performance-related dopamine differences are not driven by overt errors in monkey. **T.** Dopamine concentration changes (Δ[DA]) (right) from trials split by specific error type: all errors (top), overt fixation errors (middle) and covert omission errors (bottom). These specific rates vary throughout a single session (left), and one specific error type did not drive performance-related dopamine differences (c8c recorded on 9/24/2024).

## Discussion

We have shown that single-trial phasic dopamine signals in the primate dorsal striatum carry information about slowly evolving performance states extending over tens to hundreds of trials, simultaneously with trial-level reward value and prediction error signals. This performance-state modulation was robust, surviving regression on trial-by-trial RPE, confirming that it is not a byproduct of the prediction error signal. The relationship was temporally asymmetric, with dopamine more strongly associated with future than past performance windows, indicating a prospective link between dopamine and behavioral state. Performance-state modulation was also dissociable from behavioral and autonomic proxies of arousal — licking, saccade latency, pupil diameter, and heart rate variability — which were only weakly and inconsistently associated with performance state despite robust reward-size modulation. In the CN, performance state was a stronger modulator of dopamine than cumulative reward rate in both monkeys; this dissociation was less clear in the putamen. State-dependent dopamine signals were concentrated in the CN rather than the putamen, consistent with the oculomotor demands of the task and the known functional specialization of these regions. Together, these findings demonstrate that phasic dopamine signals in the dorsal striatum are not solely fast learning signals, but simultaneously reflect the slowly evolving internal states that shape sustained motivated performance.

The coexistence of RPE and performance-state signaling functions within the same phasic dopamine measurement was supported by converging evidence. A residuals analysis allowed us to statistically remove the portion of dopamine signal explained by trial-by-trial RPE. This analysis confirmed that performance state still modulated what remained, showing that dopamine carries separable performance-state and RPE signals. Nevertheless, the temporal structure of the two signaling functions provided an independent line of evidence: dopamine tended to be lower following a successful trial than following a failed trial (consistent with RPE encoding), whereas dopamine was higher during sustained high-performance states. That these signals run in opposite directions at adjacent timescales argues against the possibility that performance-state modulation is simply a slow reflection of accumulated prediction errors. Previous work in ventral striatum dissociated RPE from motivational signals across measurement modalities, with dopamine neuron firing in the VTA encoding RPE and local dopamine release in the nucleus accumbens tracking motivational state^53^. Our results suggest that in the primate dorsal striatum, both signals are accessible within phasic release itself. This dual-signaling function is also in line with findings that subsecond dopamine fluctuations in human caudate nucleus can carry superposed error signals about distinct reward computations^54^, and with recent work showing that mesolimbic dopamine tracks behavioral policy over hundreds of trials, dissociable from reward prediction^55^. An alternative interpretation is that apparent multiplexing of task variables in striatal dopamine can collapse to a single temporal difference error when modeled appropriately^56^. However, in our data, the performance-state signal survived explicit regression on RPE in 65 of 69 performance-modulated site-sessions, indicating that it is not reducible to the prediction error signal under any single-variable formulation.

Temporal asymmetry in dopamine responses (greater association with future than past performance) is naturally interpreted as meaning that dopamine signals the current value of task engagement^57^. If dopamine reflects how worthwhile it is to invest effort in the current task, elevated dopamine during high-performance states would correlate with upcoming performance simply because engagement states persist over many trials. Performance in our task does fluctuate on this kind of slow timescale: the behavioral autocorrelation decays over roughly 20 – 60 trials, confirming that a slow process is at work. However, this correlation admits two accounts that our data cannot fully distinguish. The first is that a slowly fluctuating internal state (motivation, arousal, or engagement) drives both dopamine release and future performance as co-consequences. Such shared signaling has precedent: slowly drifting neural population activity in primate cortex covaries with behavioral tendencies over timescales comparable to those we observe^58^. The second is that dopamine signaling is causally upstream of future performance: elevated release at striatal targets could invigorate upcoming actions or sharpen the representations guiding them. This account also has support in other contexts — activating dopamine neuron before movement onset increased the probability and vigor of future actions without triggering immediate movements^59^, and slowly ramping dopamine signals predicted the timing of upcoming self-initiated movements on individual trials^60^. These two accounts are not mutually exclusive; dopamine could both reflect an ongoing engagement state and contribute to sustaining it.

Performance-state dopamine modulation was concentrated in the CN, where effects were consistent across both monkeys and across both windowed and kernel-based analyses. This regional concentration is consistent with the behavioral demands of the task. The CN receives direct input from the frontal eye fields and contains neurons that respond during reward-guided eye movements in tasks closely related to ours^28,61–63^, making it the more task-relevant structure for our oculomotor paradigm. Performance-state modulation was also present in the putamen but was less consistent, particularly in monkey P, where only 2 of 11 sites in the putamen showed significant effects. Moreover, the dissociation from reward rate that was clear in the CN did not hold in the putamen: in monkey P, reward rate was more often associated with dopamine than performance state (14/25 vs. 5/25 site-sessions), and in monkey T, the two measures were equally prevalent. The putamen thus does not show the clean dissociation seen in the CN, which may instead reflect its weaker engagement by a task with minimal skeletomotor demands. This pattern aligns with growing evidence that dopamine signaling is spatially stratified according to the functional organization of striatal circuits^64–66^, with dopamine-gated plasticity constrained to circuits relevant to ongoing behavioral demands. One limitation qualifies this account: the CN sites we sampled from were rostral to the anterior commissure in both monkeys, whereas putamen sites were rostral in monkey P, but mostly caudal in monkey T. We therefore cannot fully dissociate whether the regional differences reflect differential engagement by performance states, the known functional specialization of CN and putamen, or the precise location of the recording sites within each structure. Tasks that place primary demands on skeletomotor output would test whether performance-state dopamine signals shift to the putamen when it becomes the more task-relevant region.

These findings raise several questions for future investigation. Establishing whether performance-state dopamine signals play a causal role in sustaining behavioral performance, as dopamine does in cue-reward learning, will require temporally precise manipulations of dopamine release during naturally occurring performance fluctuations. A second question is where these signals originate: distinct dopamine cell populations in the midbrain, or local regulatory mechanisms at striatal terminals. Multiple striatal microcircuit mechanisms can regulate dopamine release independently of midbrain spiking, including cholinergic interneuron activation of nicotinic receptors on dopamine axons^35,67^, GABAergic and autoreceptor-mediated regulation^68^, and thalamostriatal inputs that drive dopamine release through local cholinergic circuits^69^. If performance-state modulation arises partly from such local control, such as via thalamic inputs associated with arousal and attentional engagement, this could contribute to the regional specificity we observed. Complementary techniques that capture tonic or slower (>60 s) dopamine dynamics would clarify whether the performance-state signal we detect in phasic transients coexists with or is distinct from slower dopaminergic fluctuations^70,71^. Finally, the core finding replicated across both monkeys for the CN, but not as consistently in the putamen. Extending these measurements to tasks with different behavioral demands, as proposed above, would test whether performance-state dopamine signals shift to whichever striatal subregion is most task-relevant. Broader generalization to other species and behavioral contexts remains a further important goal.

## Methods

### Animals and Surgeries

Two adult female Rhesus monkeys (*Macaca mulatta*), referred to here as monkey P (∼10 kg, ∼8.5 years old at time of recording) and monkey T (∼7 kg, ∼11 years old at time of recording), were used for this study. Procedures involving monkey P were approved by the Committee on Animal Care of the Massachusetts Institute of Technology, and methods are previously reported^32,39^. Procedures involving monkey T were approved by the Institutional Animal Care and Use Committee at the University of Pittsburgh using previously reported methods^72,73^. All procedures conformed to the Guide for the Care and Use of Laboratory Animals and to the provisions of the Animal Welfare Act. Monkey P was maintained on a food-restricted schedule with liquid-food rewards delivered during recording sessions, and monkey T was maintained on a water-restricted schedule with diluted apple juice delivered during recording sessions; supplemental food or water was provided as needed to maintain target body weight and hydration. Daily health monitoring including body weight, hydration, and behavioral assessment was performed by veterinary and research staff throughout the study. During a single recording session, monkey P performed a total of 976 – 1291 trials (619 – 812 successfully) and monkey T performed a total of 1277 – 1400 trials (732 – 990 successfully). Some of the data collected for monkey P were analyzed previously^32^ and are re-analyzed here to answer a different scientific question, using different methods and task conditions, as described below.

In brief, each monkey underwent cranial chamber installation, craniotomy for brain exposure, and chronic electrode implantation. Electrodes were implanted into the caudate nucleus and putamen of the right hemisphere via form-fitting chambers and grids (Gray Matter Research); chamber implantation, craniotomy, and electrode-implantation procedures are described in detail elsewhere for monkey P^32,39^, and monkey T^72^. Carbon fiber electrodes were custom-made in the lab for application in FSCV recording of dopamine concentration changes in both monkeys^32,39,74^. Protocols for these procedures are also published online (https://doi.org/10.17504/protocols.io.kqdg32b91v25/v1, https://doi.org/10.17504/protocols.io.x54v92wd4l3e/v1, https://doi.org/10.17504/protocols.io.bp2l62m95gqe/v1).

### Behavioral Task

Both monkeys performed a visually guided reward-biased saccade task as previously described^32^. In short, each trial required the monkey to complete a saccade and fixation to an initial central cue (C) and a peripheral target (V) cue appearing to the left or the right of the C cue before receiving a large or small reward consisting of 0.1 – 0.3 mL or 1.5 – 2.8 mL liquid-food for monkey P and 90 – 100 μL or 450 – 500 μL diluted apple juice reward for monkey T. Trials in which the monkey broke fixation before reward delivery were classified as errors and were not rewarded. The association between target location and reward size (small or large) changed every 20 – 30 trials for monkey P and 45 – 75 for monkey T. Monkey T also performed a modified version of this task in which the V cue’s location-reward size association was based on its vertical position on the screen.

### Recording and Analyzing Dopamine Concentration Changes

#### Data Analysis

Data was pre-processed and prepared for analysis with custom code using standard MATLAB functions (MathWorks, MATLAB, 2025b). Analysis was performed in Python (version 3.12.3) using open-source libraries such as numpy^75^, matplotlib^76^, scipy^77^, and pandas^78^ alongside custom code.

#### Dopamine Recordings via FSCV

Dopamine recordings were obtained using FSCV as previously described^32,39,79^. Briefly, a triangular voltage waveform from –0.4 to 1.3 V (ramp rate 400 V/s) was applied to the carbon-fiber recording electrode every 100 ms (10 Hz effective sampling rate), against an Ag/AgCl reference. Dopamine oxidizes and reduces at 0.6 and –0.2 V, respectively, generating current in proportion to its concentration. The measured current were background-subtracted at the behavioral alignment event of interest (target cue) and projected onto principal components computed from *in vitro* standards of dopamine, pH, and movement artifact to estimate trial-by-trial changes in dopamine concentration (Δ[DA])^32,80^. Current signals exceeding a residual sum-of-squares tolerance *Qα* were nulled (assigned NaN in MATLAB).

#### FSCV Background Subtraction Aligned to Task Events

Background subtraction was performed by calculating a baseline value at the alignment event. This baseline was computed as the mean dopamine concentration across five time points: the sample at the event time, two samples immediately preceding, and two samples immediately following the event time. This baseline value was then subtracted from the entire trial’s dopamine recordings. Unless otherwise stated, all analyses were aligned to the second, later task event (V).

#### Computation of Single-Trial AUCs of Dopamine Concentrations

We compared Δ[DA] across trial groupings for all dopamine-focused analyses by computing trapezoidal areas under the curve (AUCs, ’cumtrapz’ in MATLAB) of the background-subtracted Δ[DA] from the task alignment event until trial completion (reward completion for monkey P and peripheral target feedback for monkey T). If a given trial had more than 30% NaN values during this time period, then the AUC for that trial was set to be NaN and excluded from statistical comparisons. Despite this exclusion from statistical comparisons, timestamps with valid recordings still contributed to trial-average visualizations.

#### Performance State Classification via Sliding Window Success Rate

We characterized trial-by-trial behavioral performance by computing a local success rate for each trial using a sliding window over the binary success sequence within each session. For a given window size *k*, the windowed success rate at trial *t* was defined as the proportion of successful trials within a window of *N* trials positioned relative to trial *t* according to a specified window location: in the start condition the window extended forward from trial *t* (i.e., trials *t* through *t + k − 1*); in the end condition it extended backward (trials *t − k + 1* through *t*); and in the middle condition the window was centered on trial *t*, extending *k*/2 (rounded down for odd *k*) trials in each direction. In all cases, a value was computed for every trial. At session edges where a full window was unavailable, the average was taken over the available (partial) window rather than treating the estimate as missing.

We classified each trial within a session into high and low performance states by applying a median split to the windowed success rates. This threshold was computed as the median of the windowed success rates evaluated only at successful trials; windowed success rates for error trials were excluded from the threshold calculation. Each trial was then labeled as “high performance” if its windowed success rate was at or above this session-specific median threshold, or “low performance” if it fell below.

#### Performance State Classification Defined by Specific Error Types

We additionally characterized trial-by-trial behavioral performance by computing a local error rate similar to the local success rate but instead using a window that counted the number of errors of a specific type that were committed in that window. The specific error types that were used in this analysis were omissions (where the animal never began fixation with the first central target) and fixation errors (where the animal began fixation with either the central or value associated target but did not hold fixation for the required time). Performance state thresholding was completed in the same way as the standard success rate method.

#### Autocorrelation Analysis of Trial Outcomes

We characterized slow fluctuations in behavioral performance state in a model-free manner by computing the trial-by-trial autocorrelation of the binary outcome sequence (success or error) on a per-session basis. If performance varied across multi-trial epochs rather than being temporally independent, positive autocorrelation should persist over multiple lags, with the rate of its decay reflecting the characteristic timescale of these fluctuations.

For each session, the binary sequence of successes and errors was mean-centered, and the autocorrelation function (ACF) was computed at lags from 0 to 50 trials. At each lag *k*, the autocovariance was obtained as

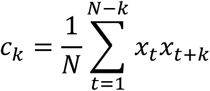

where *x_t_* denotes the mean-centered outcome on trial *t* and *N* is the total number of trials in the session. The autocorrelation at lag *k* was then classified as ρ*_k_* = *c_k_*/*c*_0_, yielding ρ_0_ = 1 by construction.

The decay of the ACF was characterized by fitting a three-parameter exponential model of the form

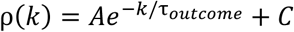

where *A* is the amplitude, τ*_outcome_* is the decay time constant (in trials) and *C* is the asymptotic baseline. The lag-0 point was excluded from the fit, as it is fixed at unity by definition and would otherwise dominate the residuals. Fits were performed by nonlinear least squares. To reduce sensitivity to local minima, the optimization was initialized form a grid of starting values for τ*_outcome_* spanning fast switches to slow drift (0.5, 2, 5, 10, 20, 50, and 100 trials); *C* was initialized to the median of the final 20 lags of the empirical ACF, and *A* to the difference between the lag-1 autocorrelation and this initial baseline. Among the resulting fits, the one yielding the highest coefficient of determination (*R*^2^) was retained as the best fit for that session. Fits with non-physical or unstable parameters were rejected by requiring *A* ≥ 0,0.3 ≤ τ*_outcome_* ≤ 500 trials, and −1 ≤ *C* ≤ 1. Sessions for which no convergent fit met these criteria were excluded from analyses and descriptions of the time constant.

### Kernel Regression of Dopamine on Outcome History

We further characterized the temporal extent over which trial outcomes covary with single-trial dopamine by fitting a basis-function kernel regression for each site-session. This analysis estimates a smooth, lag-resolved kernel relating dopamine AUC on a given trial *t* to binary trial outcomes (success or error) at lags ranging from *N* trials before to *N* trials after *t*, with past (*k* < 0) and future (*k* > 0) lags treated symmetrically.

For each site-session, we constructed a lag-design matrix *X* of size *M* × (2*N* + 1), where *M* was the number of trials with valid dopamine AUC measurements and each row contained the outcome sequence (1 for success, 0 for error) spanning *N* trials before through *N* trials after the trial on which dopamine was measured. Trials whose ±*N* window extended outside the recorded session were excluded. We projected *X* onto a basis of *K* raised-cosine bumps tiling the lag axis to enforce smoothness and reduce dimensionality following methods from prior work^81^. Basis function centers were placed at K equally spaced positions along the lag axis from −*N* to +*N*, with each bump’s half-width set to the inter-center spacing so that adjacent bumps overlapped at their half-maxima. This ensured any smoothly varying kernel could be well-approximated by their linear combination and treated all lags as a priori equivalent. We used *N* = 100, matching our sliding window size range used in subsequent analyses, and *K* = 12 throughout.

Basis coefficients, α (length *K*), were estimated by ridge regression of the mean-centered session dopamine AUC vector on the projected design matrix, *X_basis_* = *XB*. No intercept was included. The ridge penalty, λ, was selected separately for each session by five-fold cross-validation over a 25-point logarithmic grid spanning λ ∈ [10⁻², 10⁴]. The lag-resolved kernel was then reconstructed as *k* = *Bα*, yielding a (2*N* + 1)-element vector of weights over lags −*N* to +*N*. For plotting purposes, individual site-session kernels were grouped by monkey and by striatal region recorded.

### Optimal Window Definition and Determination

We defined the behavioral timescale over which dopamine most strongly covaried with performance at each recording site by sweeping across candidate window sizes and applying a two-stage significance-then-effect-size filter. The same procedure was used for both the success rate and reward rate analyses; we describe it here once in terms of success rate and note below the (minor) modifications for reward rate.

For each recording site and each of the three window positions, we swept window sizes *k* = 10 – 100. At every *k*, successful trials were partitioned into a high-performance and a low-performance group by thresholding the windowed-performance value at the session median computed on successful trials only. Performance-related dopamine differences were then assessed as described in *Performance State Classification via Sliding Window Success Rate.* This sweep produced, for each session, site, window position, and *k*, a significance flag at a pre-specified significance threshold and a raw mean difference Δ = mean_high − mean_low.

For each session, site, and window position, we then identified a single optimal window size. First, the candidate set was restricted to window sizes that reached significance at the chosen significance threshold (we used α = 0.05 unless otherwise noted). Second, among those significant sizes, the optimal window was the one maximizing the absolute raw mean difference, |Δ|. Ties (window sizes with equivalent |Δ|) were broken in favor of the smallest window size, although no analyses were done to confirm the presence of any such ties. Sites with no significant *k* had no optimal window defined, and this condition was handled on a per-analysis basis.

### Reinforcement Learning

We evaluated the extent to which each recording site’s trial-by-trial dopamine signal was consistent with this account by fitting a simple reinforcement learning model to the behavioral sequence and quantifying how strongly the resulting RPE estimates covaried with the measured dopamine AUC at each site.

We used a standard Rescorla-Wagner update rule with a single scalar value estimate *V*. *V* was initialized at 0 at the start of each session and updated on each trial according to

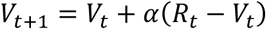

where *R_t_* is the scaled reward received on trial t and *α* ∈ (0,1] is the learning rate. The prediction error associated with trial t was defined as

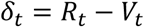

evaluated using the pre-update value estimate. The RPE sequence constructed this way was the quantity correlated against the dopamine metric.

*R_t_* was scaled such that a large reward was set to be 1 and small reward was set as the fraction of the large reward a small reward held (1/5 for monkey T and 1/14 for monkey P). This scaling preserves the relative magnitude of the task’s outcomes while placing *V*, *R*, and *RPE* on a [-1, +1] boundary.

The value update depended on what the monkey had the opportunity to observe on each trial. On successful trials, *V* was updated toward the scaled reward delivered on that trial. On error trials, the update was gated by whether the peripheral, value associated cue had been displayed before the trial ended, as determined from the recorded event markers. On error trials in which the cue was displayed, *V* was updated using *R_cue_* (the reward magnitude the monkey would have received on a successful trial under the current block contingency) inputted by forward-filling from the reward received on the next successful trial, with a correction applied if a block switched intervened between the two. On error trials in which the value associated cue was not displayed (i.e., the monkey made a fixation error during the central cue stage), *V* was held fixed and *RPE* was left undefined. This formulation makes the assumption that learning occurs only on trials in which the monkey had access to the sensory evidence required to form a reward expectation; trials ended due to error before the value associated cue’s onset carry no such evidence and are treated as non-events by the learner.

The learning rate was treated as a free parameter fit separately for each site-session combination. For each combination, we searched over *α* ∈ [0.001,1.0] for the value that maximized |*r*|, the absolute Spearman correlation between model-derived RPE and the measured dopamine AUCs across all successfully recorded trials in the session. The optimization was performed by bounded scalar minimization of the objective using Brent’s method as implemented in scipy (scipy.optimize.minimize_scalar). Although the objective maximized absolute correlation, the signed correlation coefficient to the fitted learning rate was retained for reporting such that the direction of the dopamine-RPE relationship was preserved.

We computed a session-pooled spearman correlation of dopamine to RPE for each recording site for **Figure 4c**. We pooled data for a single site using the per-session-optimized learning rates to compute RPE and used no session-based normalization of dopamine AUCs before computing the correlation.

### Residual Analysis for Dopamine-RPE Independence

We conducted an extended analysis to assess the independence of performance state-related and RPE-related dopamine signaling. For each site-session, we computed the trial-by-trial RPE using the update model previously described. We then performed standard ordinary least squares regression on our observed dopamine AUCs to computed RPE and took the residuals.

The standard analytical approach for assessing performance state-related dopamine signaling (see *Performance State Classification via Sliding Window Success Rate*) was done using the residuals to compute the test statistic and null distribution via the circular permutation method.

### Temporal Selectivity of Performance-Related Dopamine

We tested whether dopamine activity at a recording site was more strongly associated with upcoming (prospective) or recently completed (retrospective) behavioral performance by computing a per-site-session temporal selectivity bias that compared the forward- and backward-looking effect sizes at their respective optimal window sizes.

The temporal selectivity bias was computed using the effect sizes at each directions’ optimal window as:

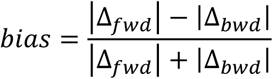

bounded on [-1, +1], where positive values indicate stronger selectivity for future performance and negative values indicate stronger selectivity for past performance. Normalizing the difference in effect size magnitude by their sum allows for across-site comparison, but this was not done in these analyses.

A bias of –1 was assigned to the site-session when there was no optimal window for future performance (i.e., there were no candidate window sizes that showed statistically significant dopamine differences between high- and low-performance states), but there was an optimal window for past performance. A bias of +1 was assigned to the site-session in the opposite case where there was an optimal window for future performance but not past performance. Site-sessions that did not show significance in either direction were labeled as incomputable for summary visualizations.

### Reward Rate Classification by Sliding Window Reward Summation

A similar local metric was computed to capture recent reward history rather than success rate. For each trial *t*, reward rate was defined as the total reward received over the *k* trials immediately preceding trial *t* (i.e., trials *t – k* through *t − 1*), with the current trial excluded. Reward amounts were scaled according to monkey-specific constants reflecting the large and small reward conditions used in the task; error trials contributed zero reward. Trials were then split into high and low reward-rate groups using the same median-split procedure described previously for success rate (see *Performance State Classification via Sliding Window Success Rate*); the threshold was taken as the median reward rate across successful trials only, and trials were assigned to high or low reward-rate groups based on their local metric.

Optimal window analyses were also performed using reward rate as the local metric to define groups as described in *Optimal Window Definition and Determination* in a verbatim manner. Because reward rate is only meaningful as a summary of recent reward history, the reward rate sweep was restricted to the ‘end’ window only.

### Comparison of Success Rate and Reward Rate on Phasic Dopamine

It is expected that monkeys would be receiving more reward whenever they are performing the behavioral task at a high success rate. However, because our task consisted of both large and small reward trials, this is not guaranteed to be the case. We computed a per-site-session bias score paralleling the temporal selectivity analysis described above with effect sizes for forward-or backward-looking windows replaced with the effect sizes for success rate and reward rate. We did this to evaluate whether our dopamine signals were more strongly modulated with previous reward history or the monkey’s performance on the task. We only presented results for forward-looking windows for the success rate component.

The procedure is structurally identical to the temporal selectivity bias computation (see *Temporal Selectivity of Performance Related Dopamine*). For each site-session combination, we drew the two optimal window sizes identified in the preceding sliding window analysis: one from the success rate optimal window sweep and one from the reward rate optimal window sweep (the latter computed over the reward history trace as described in *Reward Rate Classification by Sliding Window Reward Summation*). Using these optimal window sizes, we then determined the effect sizes for success rate (Δ*_sr_*) and reward rate (Δ*_rr_*) as also previously described.

We then applied these effect sizes to a formula similar to that used to define temporal selectivity bias,

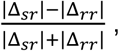

bounded again on [-1, +1] where positive values indicate that larger dopamine differences are observed for performance (success rate)-related sectioning and negative values indicate that larger dopamine differences are observed for reward-related sectioning.

Significance-based edge cases were handled as in the temporal selectivity bias analyses: site-sessions that only showed significance for success rate were assigned a score of +1, site-sessions that only showed significance for reward rate were assigned a score of –1, and site-sessions that did not show significance for either success rate or reward rate were labeled as incomputable.

### Non-Neural Physiological Measurements

Non-neural physiological signals were recorded concurrently with neural activity as proxies for internal state during task performance using methods detailed in Schwerdt et al.^32^. In brief, licking was measured as the summed absolute three-axis acceleration of the reward delivery mouthpiece via an attached accelerometer (SparkFun, MMA8452Q). Pupil diameter and eye position were measured with an infrared eye tracker (SR Research, EyeLink 1000). Pulse was recorded from an ear-clipped pulse oximeter (SparkFun, SEN-11574). All signals were routed to the electrophysiology system (Neuralynx) after attenuation, recorded synchronously with neural data, and downsampled to 1 kHz for offline analysis.

Pupil diameter was normalized on a per-recording session basis prior to analysis. For monkey T, this meant simply taking the mean and standard deviation of all recorded values while the task was underway. For monkey P however, we utilized a similar process but only included trials in which the peripheral target appeared on the right side of the screen. We took this step because pupil recordings for leftward trials were found to be unstable. These per-session summary statistics were then used to compute a z-score for pupil diameter.

### Statistics

The primary statistical test was a circular (rotation) permutation test designed to account for temporal autocorrelation in both the group label sequence and the dependent variable (e.g., performance state label classifications). A label vector was constructed by assigning each observation to one of two groups (see *Performance State Classification via Sliding Window Success Rate*), ordered by their position in the time series. The observed test statistic was the difference in group means. To generate the null distribution, the label vector was circularly shifted by *k* positions for every *k* from 1 to *N* – 1, where *N* is the length of the label vector. At each shift, the test statistic was recomputed. This procedure provides an appropriate null for testing whether two temporally structured signals are related beyond shared slow dynamics since circular rotation preserves the internal autocorrelation structure of the label sequence. Exhaustive enumeration of all *N* – 1 shifts yields a valid permutation distribution. Two-sided p-values were computed as the proportion of null statistics with absolute value greater than or equal to the observed absolute statistic.

We assessed effects pooled across multiple independent sessions in analyses containing a temporal structure using a multi-session extension of the circular permutation test. On each of 10,000 null draws, each session’s label vector was independently circularly shifted by a random amount (*k* drawn uniformly from 1 to *N_i* – 1, where *N_i* is the label vector length for session *i*). Values were then pooled across sessions separately for the shifted groups, and the pooled test statistic was computed. Draws in which any session’s shift produced an empty group were discarded, with a maximum of 50,000 total draws permitted to obtain the target 10,000 valid null statistics. P-values were computed as for the single-session test.

We used standard label-shuffle permutation tests in analyses in which no temporal structure required to be maintained in the null distribution (e.g., comparison of dopamine in large versus small reward trials). Group labels were randomly shuffled across trials 10,000 times and the difference in group means was recomputed on each shuffle to build the null distribution. Two-sided p-values were computed as the proportion of null statistics with absolute value greater than or equal to the observed statistic. Fixed random seeds were used to ensure reproducibility.

## Data Availability

Datasets used may be found through zenodo (https://doi.org/10.5281/zenodo.20419241). Protocols used in this study are available at protocols.io (dx.doi.org/10.17504/protocols.io.kqdg32b91v25/v1, dx.doi.org/10.17504/protocols.io.x54v92wd4l3e/v1, dx.doi.org/10.17504/protocols.io.bp2l62m95gqe/v1).

## Code Availability

MATLAB and Python code used to analyze data may be found at GitHub as made available through zenodo (https://doi.org/10.5281/zenodo.20617988)

## Author contributions

H.N.S., U.A., and R.M. designed and performed research. H.N.S., R.M., U.A., and J.P.H. analyzed data. H.N.S., J.P.H., and A.M.G. guided the study. All authors wrote the paper.

## Acknowledgment

The authors thank Ms. Rebecca Marflak, Dr. Julia K. Oluoch, Ms. Erica N. Griffith, Ms. Baylie S. Leveto, Ms. Stacy L. Cashman, Mr. Kevin Thiel, and animal care staff at the Systems Neuroscience Center Animal Resource Laboratory (SNARL) at the University of Pittsburgh (Pitt). This work was supported by the National Institutes of Health (NIH) (R01NS133045 to H.N.S.), Aligning Science Across Parkinson’s (ASAP-025193 to H.N.S.) through the Michael J. Fox Foundation for Parkinson’s Research (MJFF), the Saks-Kavanaugh Foundation (to A.M.G.), and the National Defense Science and Engineering Graduate (NDSEG) Fellowship Program, sponsored by the Air Force Research Laboratory (AFRL), the Office of Naval Research (ONR), and the Army Research Office (ARO) (FA9550-21-F-0003 to U.A.).

---

**Extended Data Figure 1.**
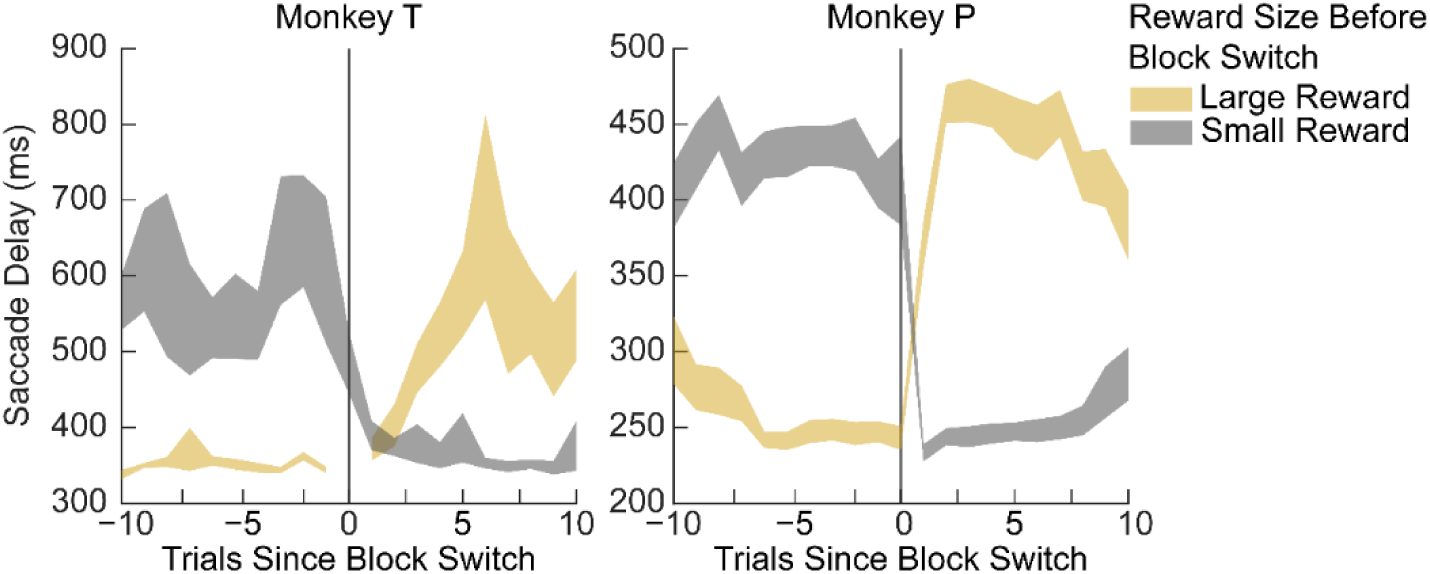
Monkeys learn the value-associated target’s location-reward contingencies quickly after a block switch. Trial-averaged saccade delays (defined as the time from when the value-associated target appears on the screen to when the monkey begins fixation on the target) around block switches when the target’s location-reward contingency switches (e.g., target on the left is associated with large reward before block switch then becomes associated with a small reward after block switch). Shading denotes ±SEM. Shading color denotes the reward size before the block switch. Both monkeys (monkey T, recorded on 3/20/2025, left; monkey P, session 94, right) show marked changes in saccade delay after a block switch such that saccade delays for targets that originally predicted a large reward increase after a block switch and saccade delays for target that originally predicted a small reward decrease after a block switch.

**Extended Data Figure 2.**
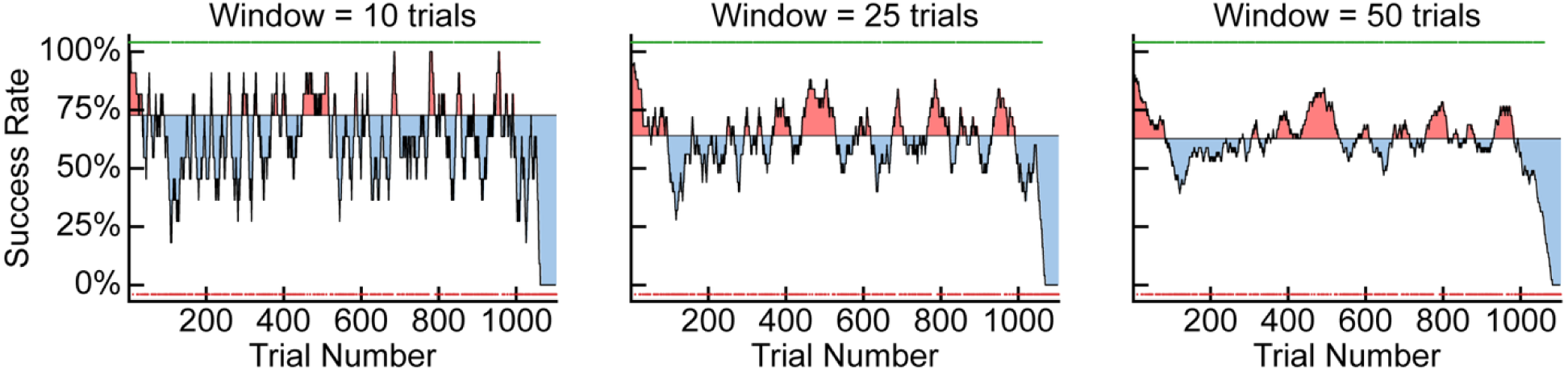
Performance state classifications arise for different window sizes within the same recording session. Moving average success rate (black) plotted as a function of trial number plotted for window lengths of 10 (left), 25 (middle) and 50 (right) trials. Shading indicates subsequent performance state classification (red: “high”, blue: “low”) while dots indicate the outcome of the corresponding trial (green: success, red: error).

**Extended Data Figure 3.**
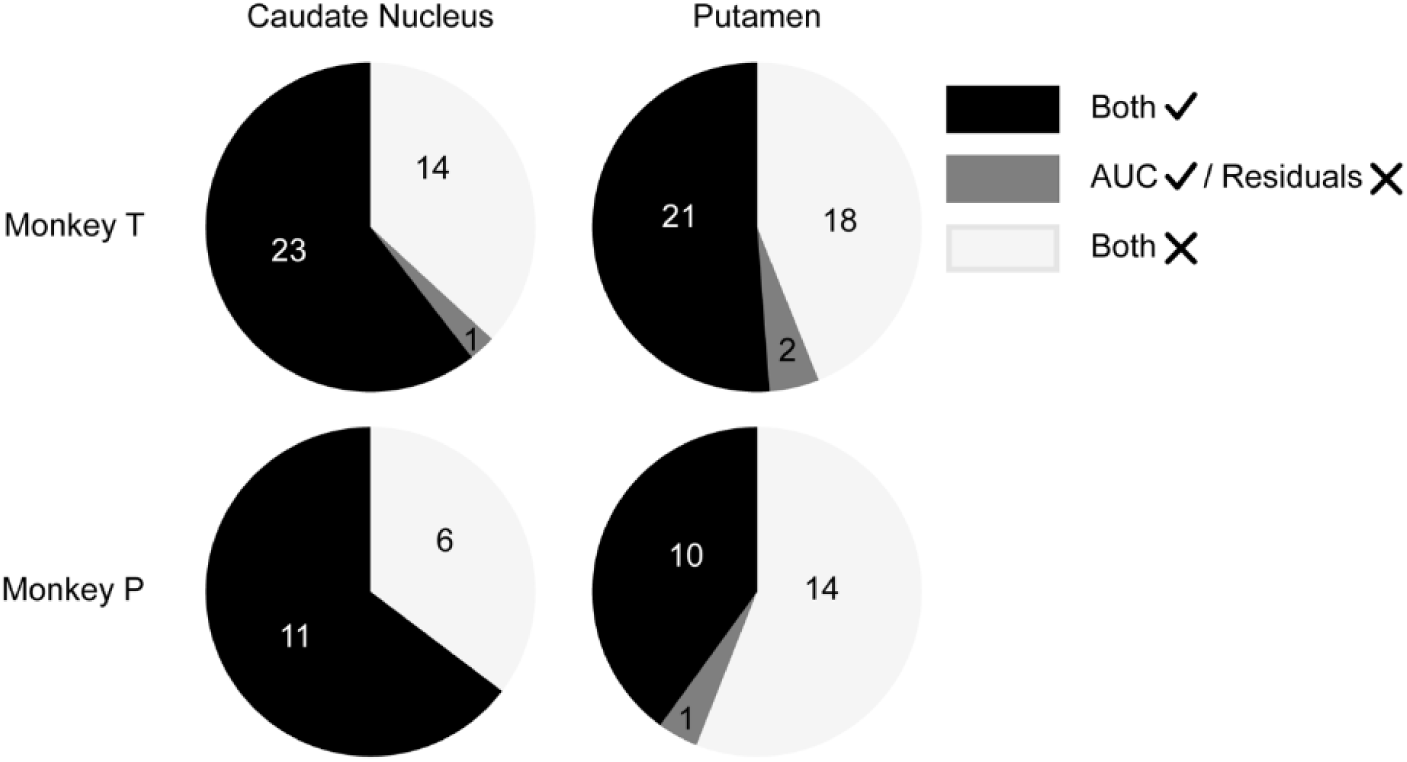
Performance state effects persist after removing contributions due to RPE. Pie charts denoting site-session counts for monkeys T (top) and P (bottom) denoting the presence of performance-phase related dopamine signaling in the caudate nucleus (left) and putamen (right). Black indicates performance-phase related dopamine signaling is present for both the raw AUC values and RPE-removed residuals. Dark gray indicates performance-related dopamine signaling is only present for raw AUC values. Light gray indicates no performance-related dopamine signaling for either raw AUC values or RPE-removed residuals.

**Extended Data Figure 4.**
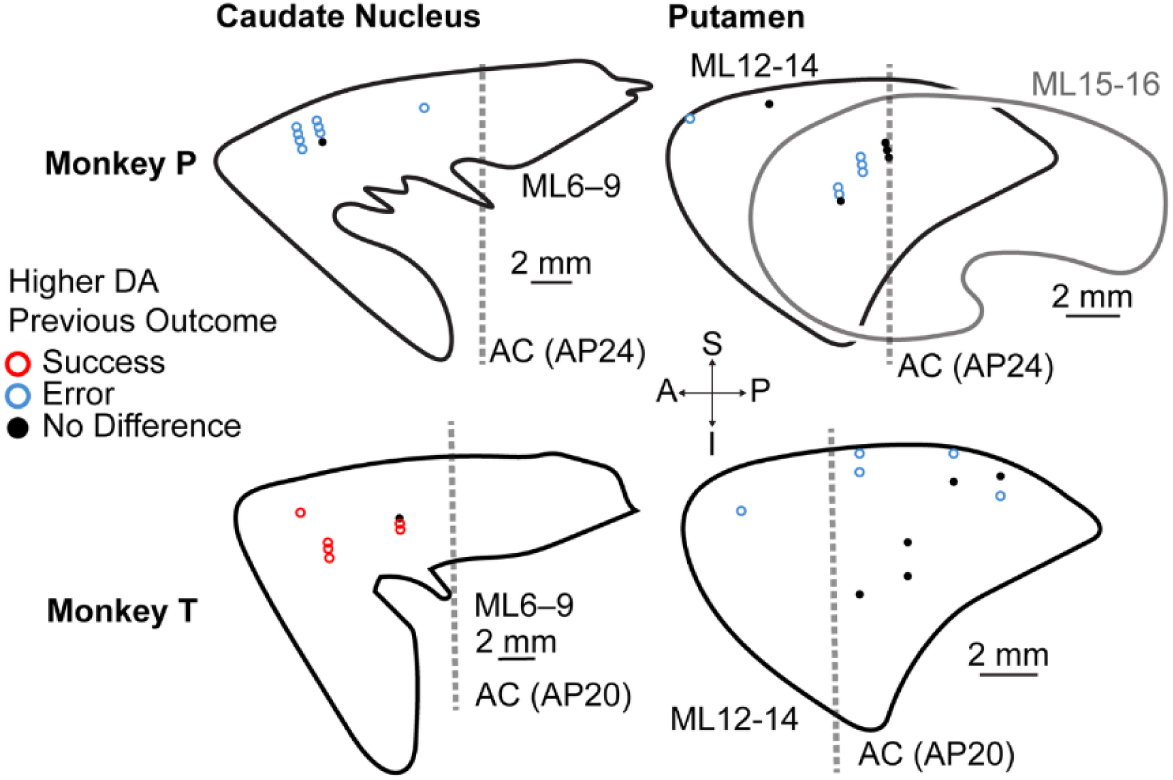
Immediate previous performance effects on current trial dopamine. Coronal sections showing recording site locations in the caudate nucleus and putamen for monkey P (top) and monkey T (bottom), color-coded by whether dopamine was greater when the previous trial was a successful or error trial

**Extended Data Figure 5.**
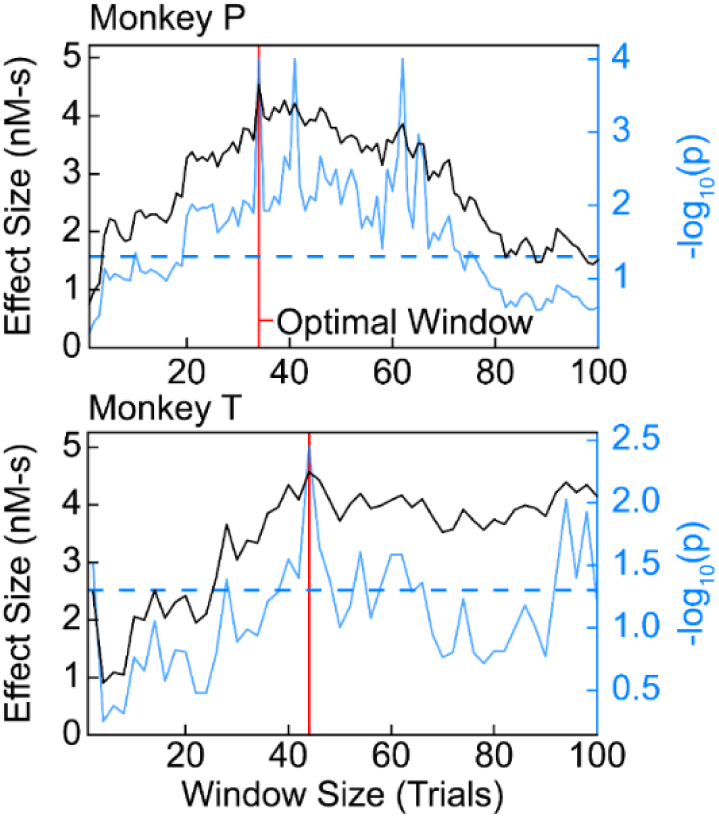
Performance state modulated dopamine responses are influenced by the duration of the performance state window. Effect size (black, see **Methods**) plotted as a function of performance state window size with p-value overlaid (blue, two-sided circular permutation test, dashed line denotes significance level of 0.05) for monkey P (top) and monkey T (bottom). Red vertical lines denote the optimal window (see **Methods**) which was used for trial-average plotting and comparative analyses (*Single-trial dopamine release is strongly modulated by prolonged performance states, Performance-state dopamine signals are stronger than reward rate modulation, especially in the CN,* see **Methods**).

## Notes

https://doi.org/10.5281/zenodo.20419241

https://doi.org/10.5281/zenodo.20617988

### Competing Interest Statement

The authors have declared no competing interest.

## References

1. Saunders, B. & More, K. R. Some habits are more work than others: Deliberate self-regulation strategy use increases with behavioral complexity, even for established habits. Journal of Personality 93, 233–246 (2025).

2. Mackworth, N. H. The Breakdown of Vigilance during Prolonged Visual Search1. Quarterly Journal of Experimental Psychology 1, 6–21 (1948).

3. Esterman, M. & Rothlein, D. Models of sustained attention. Current Opinion in Psychology 29, 174–180 (2019).

4. Salamone, J. D. & Correa, M. The Mysterious Motivational Functions of Mesolimbic Dopamine. Neuron 76, 470–485 (2012).

5. Hamid, A. A., Pettibone, J. R., Mabrouk, O. S., Hetrick, V. L., Schmidt, R., Vander Weele, C. M., Kennedy, R. T., Aragona, B. J. & Berke, J. D. Mesolimbic dopamine signals the value of work. Nat Neurosci 19, 117–126 (2016).

6. Niv, Y., Daw, N. D., Joel, D. & Dayan, P. Tonic dopamine: opportunity costs and the control of response vigor. Psychopharmacology 191, 507–520 (2006).

7. Botvinick, M. & Braver, T. Motivation and Cognitive Control: From Behavior to Neural Mechanism. Annual Review of Psychology 66, 83–113 (2015).

8. Palmiter, R. D. Dopamine Signaling in the Dorsal Striatum Is Essential for Motivated Behaviors: Lessons from Dopamine-deficient Mice. Ann N Y Acad Sci 1129, 35–46 (2008).

9. Berridge, K. C. & Robinson, T. E. What is the role of dopamine in reward: hedonic impact, reward learning, or incentive salience? Brain Res Brain Res Rev 28, 309–369 (1998).

10. Berridge, K. C. Motivation concepts in behavioral neuroscience. Physiol Behav 81, 179–209 (2004).

11. Aston-Jones, G. & Cohen, J. D. An integrative theory of locus coeruleus-norepinephrine function: adaptive gain and optimal performance. Annu Rev Neurosci 28, 403–450 (2005).

12. Flavell, S. W., Gogolla, N., Lovett-Barron, M. & Zelikowsky, M. The emergence and influence of internal states. Neuron 110, 2545–2570 (2022).

13. Calhoun, A. J., Pillow, J. W. & Murthy, M. Unsupervised identification of the internal states that shape natural behavior. Nat Neurosci 22, 2040–2049 (2019).

14. Zhang, S. X., Lutas, A., Yang, S., Diaz, A., Fluhr, H., Nagel, G., Gao, S. & Andermann, M. L. Hypothalamic dopamine neurons motivate mating through persistent cAMP signalling. Nature 597, 245–249 (2021).

15. Pessiglione, M., Vinckier, F., Bouret, S., Daunizeau, J. & Le Bouc, R. Why not try harder? Computational approach to motivation deficits in neuro-psychiatric diseases. Brain 141, 629–650 (2018).

16. Bromberg-Martin, E. S., Matsumoto, M. & Hikosaka, O. Dopamine in Motivational Control: Rewarding, Aversive, and Alerting. Neuron 68, 815–834 (2010).

17. Schultz, W. Predictive Reward Signal of Dopamine Neurons. Journal of Neurophysiology 80, 1–27 (1998).

18. Wise, R. A. Dopamine, learning and motivation. Nat Rev Neurosci 5, 483–494 (2004).

19. Schultz, W., Dayan, P. & Montague, P. R. A neural substrate of prediction and reward. Science 275, 1593–1599 (1997).

20. Fiorillo, C. D., Tobler, P. N. & Schultz, W. Discrete Coding of Reward Probability and Uncertainty by Dopamine Neurons. Science 299, 1898–1902 (2003).

21. Bayer, H. M. & Glimcher, P. W. Midbrain Dopamine Neurons Encode a Quantitative Reward Prediction Error Signal. Neuron 47, 129–141 (2005).

22. Matsumoto, M. & Hikosaka, O. Two types of dopamine neuron distinctly convey positive and negative motivational signals. Nature 459, 837–841 (2009).

23. Cohen, J. Y., Haesler, S., Vong, L., Lowell, B. B. & Uchida, N. Neuron-type-specific signals for reward and punishment in the ventral tegmental area. Nature 482, 85–88 (2012).

24. Starkweather, C. K., Babayan, B. M., Uchida, N. & Gershman, S. J. Dopamine reward prediction errors reflect hidden-state inference across time. Nat Neurosci 20, 581–589 (2017).

25. Stuber, G. D., Klanker, M., de Ridder, B., Bowers, M. S., Joosten, R. N., Feenstra, M. G. & Bonci, A. Reward-predictive cues enhance excitatory synaptic strength onto midbrain dopamine neurons. Science 321, 1690–1692 (2008).

26. Grove, J. C. R., Gray, L. A., La Santa Medina, N., Sivakumar, N., Ahn, J. S., Corpuz, T. V., Berke, J. D., Kreitzer, A. C. & Knight, Z. A. Dopamine subsystems that track internal states. Nature 608, 374–380 (2022).

27. Mohebi, A., Wei, W., Pelattini, L., Kim, K. & Berke, J. D. Dopamine transients follow a striatal gradient of reward time horizons. Nat Neurosci 27, 737–746 (2024).

28. Kawagoe, R., Takikawa, Y. & Hikosaka, O. Reward-Predicting Activity of Dopamine and Caudate Neurons—A Possible Mechanism of Motivational Control of Saccadic Eye Movement. Journal of Neurophysiology 91, 1013–1024 (2004).

29. Kawagoe, R., Takikawa, Y. & Hikosaka, O. Expectation of reward modulates cognitive signals in the basal ganglia. Nat Neurosci 1, 411–416 (1998).

30. Hikosaka, O., Takikawa, Y. & Kawagoe, R. Role of the Basal Ganglia in the Control of Purposive Saccadic Eye Movements. Physiological Reviews 80, 953–978 (2000).

31. Graybiel, A. M. Habits, Rituals, and the Evaluative Brain. Annu. Rev. Neurosci. 31, 359–387 (2008).

32. Schwerdt, H. N., Amemori, K. & Graybiel, A. M. Dopamine and beta-band oscillations differentially link to striatal value and motor control. Science Advances https://www.science.org/doi/10.1126/sciadv.abb9226 (2020).

33. van Elzelingen, W., Warnaar, P., Matos, J., Bastet, W., Jonkman, R., Smulders, D., Goedhoop, J., Denys, D., Arbab, T. & Willuhn, I. Striatal dopamine signals are region specific and temporally stable across action-sequence habit formation. Current Biology 32, 1163–1174.e6 (2022).

34. Hamid, A. A., Frank, M. J. & Moore, C. I. Wave-like dopamine dynamics as a mechanism for spatiotemporal credit assignment. Cell 184, 2733–2749.e16 (2021).

35. Threlfell, S., Lalic, T., Platt, N. J., Jennings, K. A., Deisseroth, K. & Cragg, S. J. Striatal dopamine release is triggered by synchronized activity in cholinergic interneurons. Neuron 75, 58–64 (2012).

36. Lazaridis, I., Ahn, G., Hirokane, K., Choi, W. & Graybiel, A. M. Striatal Astrocytes Influence Dopamine Dynamics and Behavioral State Transitions. 2024.12.01.626240 Preprint at 10.1101/2024.12.01.626240 (2024).

37. Keithley, R. B., Heien, M. L. & Wightman, R. M. Multivariate concentration determination using principal component regression with residual analysis. Trends Analyt Chem 28, 1127–1136 (2009).

38. Schwerdt, H. N., Zhang, E., Kim, M. J., Yoshida, T., Stanwicks, L., Amemori, S., Dagdeviren, H. E., Langer, R., Cima, M. J. & Graybiel, A. M. Cellular-scale probes enable stable chronic subsecond monitoring of dopamine neurochemicals in a rodent model. Commun Biol 1, 144 (2018).

39. Schwerdt, H. N., Shimazu, H., Amemori, K., Amemori, S., Tierney, P. L., Gibson, D. J., Hong, S., Yoshida, T., Langer, R., Cima, M. J. & Graybiel, A. M. Long-term dopamine neurochemical monitoring in primates. Proceedings of the National Academy of Sciences 114, 13260–13265 (2017).

40. Schwerdt, H. N., Shimazu, H., Amemori, K., Amemori, S., Tierney, P. L., Gibson, D. J., Hong, S., Yoshida, T., Langer, R., Cima, M. J. & Graybiel, A. M. Long-term dopamine neurochemical monitoring in primates. Proceedings of the National Academy of Sciences 114, 13260–13265 (2017).

41. Choi, J., Amjad, U., Murray, R., Shrivastav, R., Teichert, T., Goodell, B., Olson, M., Schaeffer, D. J., Oluoch, J. K. & Schwerdt, H. N. Aseptic, semi-sealed cranial chamber implants for chronic multi-channel neurochemical and electrophysiological neural recording in nonhuman primates. Journal of Neuroscience Methods 420, 110467 (2025).

42. Patriarchi, T., Cho, J. R., Merten, K., Howe, M. W., Marley, A., Xiong, W.-H., Folk, R. W., Broussard, G. J., Liang, R., Jang, M. J., Zhong, H., Dombeck, D., von Zastrow, M., Nimmerjahn, A., Gradinaru, V., Williams, J. T. & Tian, L. Ultrafast neuronal imaging of dopamine dynamics with designed genetically encoded sensors. Science 360, eaat4422 (2018).

43. Cromwell, H. C. & Schultz, W. Effects of Expectations for Different Reward Magnitudes on Neuronal Activity in Primate Striatum. Journal of Neurophysiology 89, 2823–2838 (2003).

44. Joshi, S., Li, Y., Kalwani, R. M. & Gold, J. I. Relationships between Pupil Diameter and Neuronal Activity in the Locus Coeruleus, Colliculi, and Cingulate Cortex. Neuron 89, 221–234 (2016).

45. Berntson, G. G., Thomas Bigger Jr., J., Eckberg, D. L., Grossman, P., Kaufmann, P. G., Malik, M., Nagaraja, H. N., Porges, S. W., Saul, J. P., Stone, P. H. & Van Der Molen, M. W. Heart rate variability: Origins, methods, and interpretive caveats. Psychophysiology 34, 623–648 (1997).

46. Thayer, J. F., Hansen, A. L., Saus-Rose, E. & Johnsen, B. H. Heart Rate Variability, Prefrontal Neural Function, and Cognitive Performance: The Neurovisceral Integration Perspective on Self-regulation, Adaptation, and Health. Ann Behav Med 37, 141–153 (2009).

47. Taghia, J., Cai, W., Ryali, S., Kochalka, J., Nicholas, J., Chen, T. & Menon, V. Uncovering hidden brain state dynamics that regulate performance and decision-making during cognition. Nat Commun 9, 2505 (2018).

48. Steinberg, E. E., Keiflin, R., Boivin, J. R., Witten, I. B., Deisseroth, K. & Janak, P. H. A causal link between prediction errors, dopamine neurons and learning. Nat Neurosci 16, 966–973 (2013).

49. Tsai, H.-C., Zhang, F., Adamantidis, A., Stuber, G. D., Bonci, A., de Lecea, L. & Deisseroth, K. Phasic Firing in Dopaminergic Neurons Is Sufficient for Behavioral Conditioning. Science 324, 1080–1084 (2009).

50. Dabney, W., Kurth-Nelson, Z., Uchida, N., Starkweather, C. K., Hassabis, D., Munos, R. & Botvinick, M. A distributional code for value in dopamine-based reinforcement learning. Nature 577, 671–675 (2020).

51. Lee, D., Seo, H. & Jung, M. W. Neural Basis of Reinforcement Learning and Decision Making. Annu. Rev. Neurosci. 35, 287–308 (2012).

52. Tobler, P. N., Fiorillo, C. D. & Schultz, W. Adaptive Coding of Reward Value by Dopamine Neurons. Science 307, 1642–1645 (2005).

53. Mohebi, A., Pettibone, J. R., Hamid, A. A., Wong, J.-M. T., Vinson, L. T., Patriarchi, T., Tian, L., Kennedy, R. T. & Berke, J. D. Dissociable dopamine dynamics for learning and motivation. Nature 570, 65–70 (2019).

54. Kishida, K. T., Saez, I., Lohrenz, T., Witcher, M. R., Laxton, A. W., Tatter, S. B., White, J. P., Ellis, T. L., Phillips, P. E. M. & Montague, P. R. Subsecond dopamine fluctuations in human striatum encode superposed error signals about actual and counterfactual reward. Proceedings of the National Academy of Sciences 113, 200–205 (2016).

55. Coddington, L. T., Lindo, S. E. & Dudman, J. T. Mesolimbic dopamine adapts the rate of learning from action. Nature 614, 294–302 (2023).

56. Tsutsui-Kimura, I., Matsumoto, H., Akiti, K., Yamada, M. M., Uchida, N. & Watabe-Uchida, M. Distinct temporal difference error signals in dopamine axons in three regions of the striatum in a decision-making task. eLife 9, e62390 (2020).

57. Berke, J. D. What does dopamine mean? Nat Neurosci 21, 787–793 (2018).

58. Cowley, B. R., Snyder, A. C., Acar, K., Williamson, R. C., Yu, B. M. & Smith, M. A. Slow Drift of Neural Activity as a Signature of Impulsivity in Macaque Visual and Prefrontal Cortex. Neuron 108, 551–567.e8 (2020).

59. da Silva, J. A., Tecuapetla, F., Paixão, V. & Costa, R. M. Dopamine neuron activity before action initiation gates and invigorates future movements. Nature 554, 244–248 (2018).

60. Hamilos, A. E., Spedicato, G., Hong, Y., Sun, F., Li, Y. & Assad, J. A. Slowly evolving dopaminergic activity modulates the moment-to-moment probability of reward-related self-timed movements. eLife 10, e62583 (2021).

61. Hikosaka, O., Sakamoto, M. & Usui, S. Functional properties of monkey caudate neurons. I. Activities related to saccadic eye movements. Journal of Neurophysiology 61, 780–798 (1989).

62. Hikosaka, O., Sakamoto, M. & Usui, S. Functional properties of monkey caudate neurons. III. Activities related to expectation of target and reward. Journal of Neurophysiology 61, 814–832 (1989).

63. Nakamura, K. & Hikosaka, O. Role of Dopamine in the Primate Caudate Nucleus in Reward Modulation of Saccades. J. Neurosci. 26, 5360–5369 (2006).

64. Reynolds, J. N. J., Hyland, B. I. & Wickens, J. R. A cellular mechanism of reward-related learning. Nature 413, 67–70 (2001).

65. Yagishita, S., Hayashi-Takagi, A., Ellis-Davies, G. C. R., Urakubo, H., Ishii, S. & Kasai, H. A critical time window for dopamine actions on the structural plasticity of dendritic spines. Science 345, 1616–1620 (2014).

66. Gerstner, W., Lehmann, M., Liakoni, V., Corneil, D. & Brea, J. Eligibility Traces and Plasticity on Behavioral Time Scales: Experimental Support of NeoHebbian Three-Factor Learning Rules. Front. Neural Circuits 12, (2018).

67. Cachope, R., Mateo, Y., Mathur, B. N., Irving, J., Wang, H.-L., Morales, M., Lovinger, D. M. & Cheer, J. F. Selective Activation of Cholinergic Interneurons Enhances Accumbal Phasic Dopamine Release: Setting the Tone for Reward Processing. Cell Reports 2, 33–41 (2012).

68. Sulzer, D., Cragg, S. J. & Rice, M. E. Striatal dopamine neurotransmission: Regulation of release and uptake. Basal Ganglia 6, 123–148 (2016).

69. Cover, K. K., Gyawali, U., Kerkhoff, W. G., Patton, M. H., Mu, C., White, M. G., Marquardt, A. E., Roberts, B. M., Cheer, J. F. & Mathur, B. N. Activation of the Rostral Intralaminar Thalamus Drives Reinforcement through Striatal Dopamine Release. Cell Reports 26, 1389–1398.e3 (2019).

70. Atcherley, C. W., Wood, K. M., Parent, K. L., Hashemi, P. & Heien, M. L. The coaction of tonic and phasic dopamine dynamics. *Chemical communications (Cambridge*, England) 51, 2235 (2015).

71. Movassaghi, C. S., Perrotta, K. A., Yang, H., Iyer, R., Cheng, X., Dagher, M., Fillol, M. A. & Andrews, A. M. Simultaneous serotonin and dopamine monitoring across timescales by rapid pulse voltammetry with partial least squares regression. Anal Bioanal Chem 413, 6747–6767 (2021).

72. Choi, J., Amjad, U., Murray, R., Shrivastav, R., Teichert, T., Goodell, B., Olson, M., Schaeffer, D. J., Oluoch, J. K. & Schwerdt, H. N. Aseptic, semi-sealed cranial chamber implants for chronic multi-channel neurochemical and electrophysiological neural recording in nonhuman primates. Journal of Neuroscience Methods 420, 110467 (2025).

73. Amjad, U., Mahajan, S., Choi, J., Shrivastav, R., Murray, R., Somich, A., Coyne, O. & Schwerdt, H. N. Microinvasive Probes for Monitoring Electrical and Chemical Neural Activity in Nonhuman Primates. ACS Chem. Neurosci. 16, 2237–2247 (2025).

74. Schwerdt, H. N., Jung Kim, M., Amemori, S., Homma, D., Yoshida, T., Shimazu, H., Yerramreddy, H., Karasan, E., Langer, R., M. Graybiel, A. & J. Cima, M. Subcellular probes for neurochemical recording from multiple brain sites. Lab on a Chip 17, 1104–1115 (2017).

75. Harris, C. R., Millman, K. J., van der Walt, S. J., Gommers, R., Virtanen, P., Cournapeau, D., Wieser, E., Taylor, J., Berg, S., Smith, N. J., Kern, R., Picus, M., Hoyer, S., van Kerkwijk, M. H., Brett, M., Haldane, A., del Río, J. F., Wiebe, M., Peterson, P., Gérard-Marchant, P., Sheppard, K., Reddy, T., Weckesser, W., Abbasi, H., Gohlke, C. & Oliphant, T. E. Array programming with NumPy. Nature 585, 357–362 (2020).

76. Hunter, J. D. Matplotlib: A 2D Graphics Environment. Computing in Science & Engineering 9, 90–95 (2007).

77. Virtanen, P., Gommers, R., Oliphant, T. E., Haberland, M., Reddy, T., Cournapeau, D., Burovski, E., Peterson, P., Weckesser, W., Bright, J., van der Walt, S. J., Brett, M., Wilson, J., Millman, K. J., Mayorov, N., Nelson, A. R. J., Jones, E., Kern, R., Larson, E., Carey, C. J., Polat, İ., Feng, Y., Moore, E. W., VanderPlas, J., Laxalde, D., Perktold, J., Cimrman, R., Henriksen, I., Quintero, E. A., Harris, C. R., Archibald, A. M., Ribeiro, A. H., Pedregosa, F. & van Mulbregt, P. SciPy 1.0: fundamental algorithms for scientific computing in Python. Nat Methods 17, 261–272 (2020).

78. McKinney, W. Data Structures for Statistical Computing in Python. in 56–61 (Austin, Texas, 2010). doi:10.25080/Majora-92bf1922-00a.

79. Amjad, U., Choi, J., Gibson, D. J., Murray, R., Graybiel, A. M. & Schwerdt, H. N. Synchronous Measurements of Extracellular Action Potentials and Neurochemical Activity with Carbon Fiber Electrodes in Nonhuman Primates. eNeuro 11, (2024).

80. Keithley, R. B. & Wightman, R. M. Assessing Principal Component Regression Prediction of Neurochemicals Detected with Fast-Scan Cyclic Voltammetry. ACS Chem. Neurosci. 2, 514–525 (2011).

81. Pillow, J. W., Shlens, J., Paninski, L., Sher, A., Litke, A. M., Chichilnisky, E. J. & Simoncelli, E. P. Spatio-temporal correlations and visual signalling in a complete neuronal population. Nature 454, 995–999 (2008).

